# Variation in the Microbiomes of the Basidiomycete Fungi *Scleroderma citrinum* (Pers.) and *Pisolithus arhizus* (Pers.): a tale of two saprotrophs

**DOI:** 10.1101/2023.07.25.550551

**Authors:** Ken Cullings, Shilpa R. Bhardwaj, Michael Spector

## Abstract

In this study we used high throughput DNA sequencing and ICP-MS to compare the microbiome of the common earthball fungus, *Scleroderma citrinum* (Pers.) to that of its sister taxon in the Sclerodermataceae, *Pisolithus arhizus* (Scop.). ICP-MS analysis demonstrates that *S. citrinum* is enriched in silica, sulfur and zinc relative to *P. arhizus*, while *P. arhizus* is enriched in arsenic, calcium, cadmium, cobalt, copper, lithium, magnesium, molybdenum, nickel, potassium and vanadium. Statistical analysis of molecular data indicates that the microbiome of *P. arhizus* is both richer and more diverse than that of *S. citrinum*, and that the microbiomes are significantly different with that of *S. citrinum* being enriched in Cyanobacteria represented by the chloroplast of a photosynthetic, cryptoendolithic red alga, Saccharibacteria (TM-7), and Planctomycetes, while that of *P. arhizus* is enriched in Gemmatimonadetes, Latescibacteria, Elusomicrobia, and Tectomicrobia. Further, the *P. arhizus* microbiome is enriched in anaerobes relatives to that of *S. citrinum*, probably reflecting anaerobic zones previously measured in *P. arhizus*. Together, the data indicate diverse microbiomes comprised of aromatic hydrocarbon-degrading, metal- and radiotolerant bacteria, indicating that these fungi may provide a rich source of novel microbes suitable for bioremediation strategies.

## Introduction

The study of microbiomes is critical to our understanding of organismal and ecosystem evolution and functioning. The advent of photosynthetic microbes and symbioses between photosynthetic and non-photosynthetic microbes changed the planet from one devoid of oxygen to an aerobic world that supports the vast diversity of life we know today (e.g., reviewed by Blaser et al. 2016). Microbes power critical ecosystem functions such as carbon, phosphorus and nitrogen cycling, and through symbioses with eukaryotes provide an array of services from geochemical cycling to pathogen resistance to ecosystem sustainability in the face of anthropomorphic change (Blaset et al. 2016; Dubilier et al. 2015). Symbioses between microbes and eukaryotes allow higher organisms to live and thrive in inhospitable habitats such as deep sea hydrothermal vents (e.g., Cavanaugh et al. 1981), allow both invertebrate and vertebrate animals access to substrates such as lignin and cellulose that would otherwise defy processing (E.g., Breznak 1982; Ceja-Navarro et al. 2015; Geib et al. 2008) and aid plants in growth, uptake and fixation of nitrogen, stress tolerance, and pathogen protection (reviewed by Schlaeppi and Bulgarelli 2015; Turner et al. 2013).

In our work, we focus on the microbiomes of basidiomycete fungi, the mushroom formers. The study of mushroom microbiomes is a nascent field with modern high throughput metagenomics methods providing new insight into their structure, evolution and function at a rapid pace. Recent work shows that mushrooms play host to a variety of microbes, including Archaea, Proteobacteria, Bacteriodetes, Firmicutes, and Actinobacteria (e.g., Barbieri et al. 2005; Benucci and Bonito 2016; Pent et al. 2018; Pent et al. 2020; Rinta-Kanto et al. 2018).

These bacteria can impact nitrogen nutrition and gene expression in their hosts, influence spore dispersal, aid in mycelial growth and fruiting body formation, and also act as biocontrol agents through mycotoxin production (e.g., Bahram et al. 2018; Barbieri et al. 2010; Cho et al. 2003; Deveau et al. 2018; Lackner et al. 2009; Torres-Cortes et al. 2015). Factors that influence and control the composition of these microbial communities are only now coming to light. It has been hypothesized that host genetics, pH, C/N physiology, fruiting body type and mode of nutrition could all influence microbiome community structure in fungi (e.g., Barros et al. 2007; Barros et al. 2008; Pent et al. 2018; Pent et al. 2020; Rinta-Kanto et al. 2018). Evidence for the latter relationships comes in good part from comparisons of saprophytes (fungi that obtain carbon via enzymatic breakdown of soil substrates) and ectomycorrhizae (fungi that receive carbon via symbioses with tree roots); data from these studies suggest that fruiting bodies of saprophytic basidiomycetes can be dominated by Actinobacteria, the *Burkholderia*-*Caballeronia*-*Paraburkholderia* complex of the Betaproteobacteria, Acidobacteria and Verrucobacteria (e.g., (Barbier et al. 2005; Benucci and Bonito 2016; Pent et al. 2018; Pent et al. 2020; Quandt et al. 2015; Zagriadskaia et al. 2013), while these taxa can be rare in ectomycorrhizal strains of fungi. Indeed, our previous work with *P. arhizus* in Yellowstone demonstrated that many of these taxa are prevalent in this fungus, which though often forms ectomycorrhizal associations, does not do so in Yellowstone thermal soils (Cullings and Makhija, Cullings et al. 2020).

In this study, we used high throughput DNA sequencing methods (itags) to investigate the microbiome of the basidiomycete fungus *Scleroderma citrinum* (Pers.). *Sclerdoerma* is a member of the basidiomycete family, Sclerodermataceae, the “common earthballs” (Figure 1), a group that is characterized by closed fruiting structures. The family has its origins in the Cretaceous, and is widely distributed globally being present in both tropical and temperate habitats (reviewed by Wilson et al. 2012). *Scleroderma citrinum*, the focus of this study, is a fungal extremophile that can tolerate elevated temperatures and dry conditions and occupies habitats ranging from mine heaps, to geothermal soils, to Antarctic lakes, to ore roasting beds (e.g., Mrak et al. 2017). *Scleroderma* strains can tolerate high levels of heavy metal contamination (e.g., Agganan et al. 2015; Mleczko 2004), accumulate and concentrate toxic elements, notably Zn and S (e.g., Medve and Sayre 1994), and thrive in sites contaminated with high levels of Cu, Pb, Zn and Cd (e.g., Rudawska et al. 2011; Silva et al. 2013; Roccotiello et al. 2015). *Scleroderma* species are often ectomycorrhizal (ECM) and when functioning as such can enhance metal tolerance in their ECM hosts (e.g., Jones and Hutchinson 1988; Carrillo-González and González-Chávez 2012). However, *Scleroderma* strains can also be saprotrophic depending upon site conditions (e.g., Jeffries 1999; Mrak et al. 2017). Indeed, in our previous survey of the ectomycorrhizal community of the field site utilized in this study, we found that both *S. citrinum* and a sister taxon (*Pisolithus arhizus*) were either not present in the ECM community or present in such low frequency as to go undetected. (Cullings and Makhija 2001). Rather, our data showed that the ECM community of pines growing in this thermal area was dominated by species of *Democybe* and *Inocybe*, both agaricoid (gilled) basidiomycete fungi.

**Figure 1:**
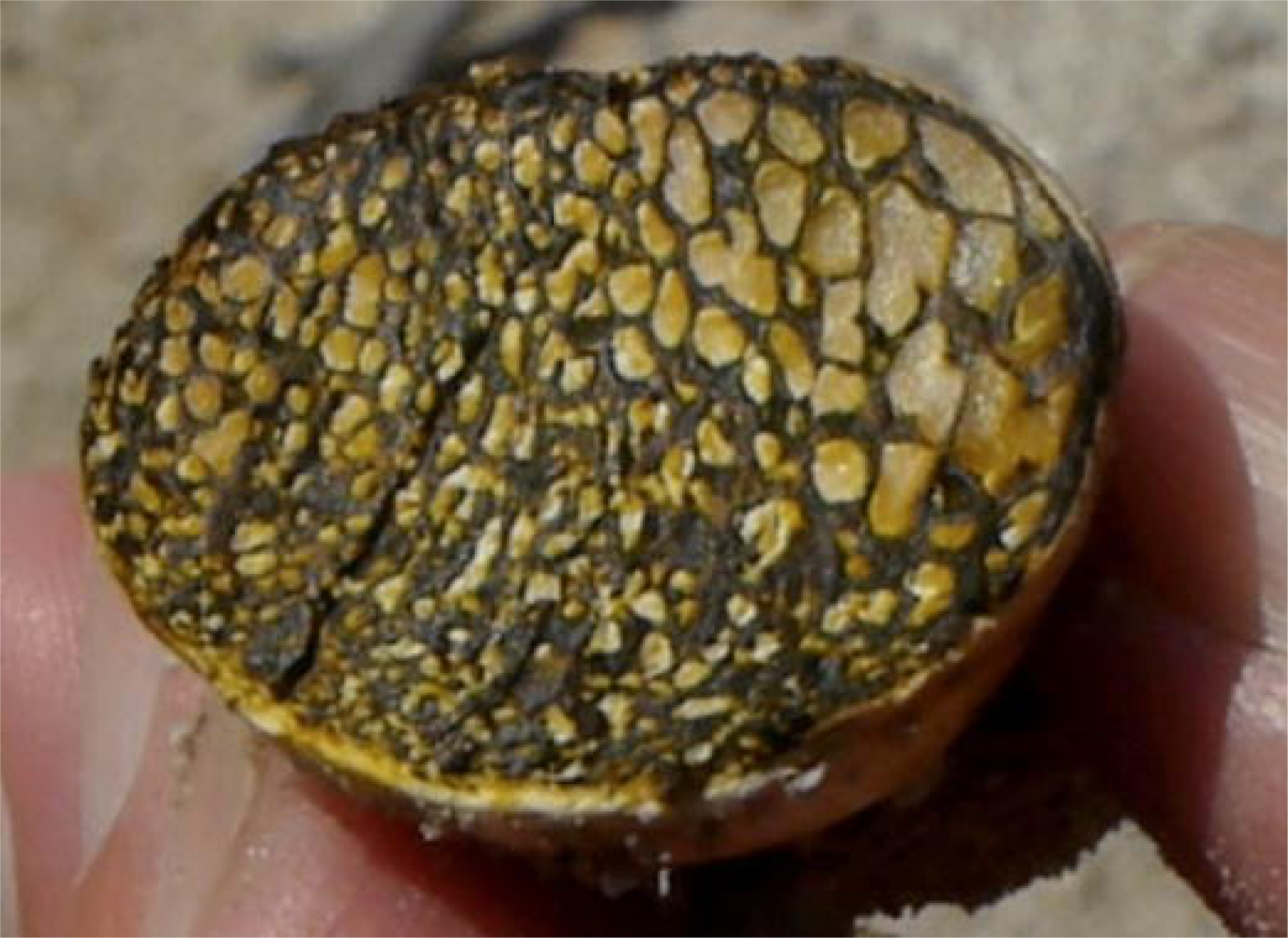
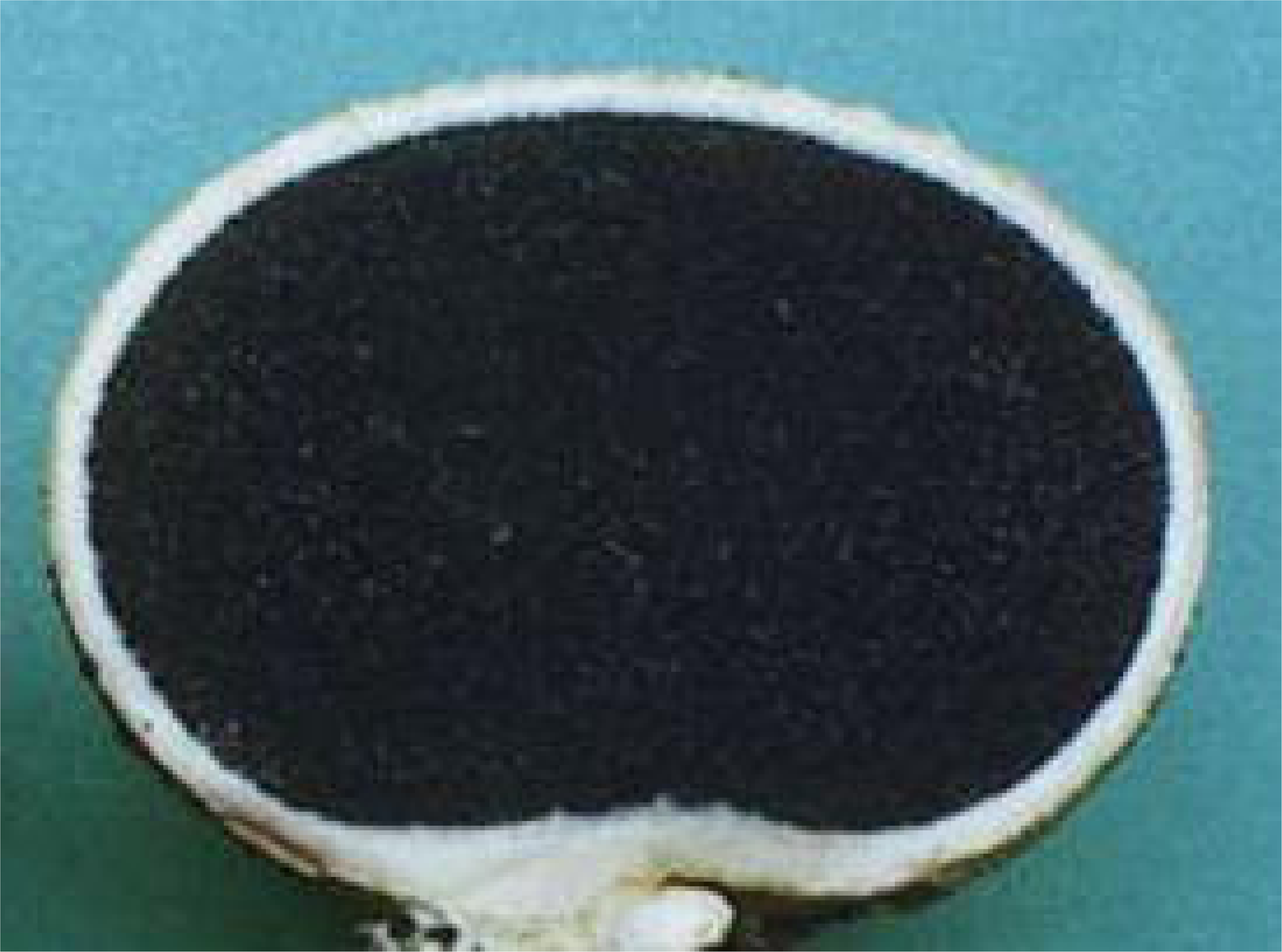
Photographs of the gleba of a) *S. citrinum* and b) *P. arhizus*. Granular inclusions within *P. arhizus* are comprised of pure silica coated in sulfur (Cullings et al 2020).

Both *S. citrinum* and *P. arhizus* form closed fruiting bodies (Figure 1) and inhabit thermal soils in Yellowstone National Park (Cullings and Makhija 2001; Cullings et al. 2020). While the two species are similar in outward appearance there are obvious, visual differences between in their internal structure (Figure 1); the gleba of *P. arhizus* contains visible inclusions that are comprised of granular Si coated in elemental S (Cullings et al. 2020), while these inclusions are conspicuously absent from the gleba of *S. citrinum*. Our data also indicated that in addition to Si and S, the fruiting bodies of *P. arhizus* concentrate Cu, Mg, Mn, Ni and Zn relative to soils. Though no direct comparisons between these two species have as yet been performed, the work of others has shown that *S. citrinum* accumulates Po, Pb (Szymańska and Strumińska-Parulska 2020). In our previous study, we demonstrated that the microbiome of *P. arhizus* shelters a rich microbiome that is comprised of metal-tolerant, hydrocarbon-degrading and chemosynthetic bacteria, photosynthetic algae, and also two potential new phyla in the Candidate Phylum Radiation (CPR) (Cullings et al. 2020). In this study, we used combined elemental and molecular datasets to a) more fully characterize the elemental components of the *S. citrinum* fruiting body and compare to that of *P. arhizus*, and b) to test the hypothesis that the diversity of the *S. citrinum* microbiome would be significantly different from that of *P. arhizus* individuals.

## Methods

### Sample Collection

This study was conducted at the Norris Annex, adjacent to Norris Geyser Basin in Yellowstone Park at the following GPS coordinates; Site 1: Lat: 44.711488

Long: -110.552502, Site 2: Lat: 44.731004, Long: -110.699787, Site 3: Lat: 44.740515 Long: -110.699830. These are the same sites used in our previous *P. arhizus* studies (Cullings and Makhija 2001, Cullings et al. 2020) and the full site characteristics are described in the former.

We collected nine pairs of *S. citrinum* and *P. arhizus* samples, with individuals of each species in the pair occurring within a meter of each other. All *S. citrinum* and *P. arhizus* sporocarps were collected aseptically into sterile plastic bags and placed immediately on dry ice for transport and storage at -80°C.

### Elemental Analysis

We performed a quantitative ICP-MS analysis on the nine pairs of *P. arhizus S. citrinum* individuals for a total of 18 analyses. Analysis was performed by GNS Science, in Taupo, New Zealand. Fungal tissue from each of the 18 individuals (approximately 1 g) was excised and weighed into a Teflon beaker, with all weights recorded to 3 decimal points. To the beaker, 4.0 mL of 1:1 HNO_3_ (trace-metal grade A509; Fisher Scientific) and 10.0 mL of 1:4 HCl (trace grade H1196; Fisher Scientific) was added and beakers were covered with a watch glass. The fungal material was then digested at 90 – 95 °C. Samples were then refluxed for 30 minutes, then allowed to cool at room temperature. The digested samples were cooled then transferred to a 100mL volumetric flask and diluted to 100 mL with de-ionized H_2_O for elemental analysis. Then, 20 mL of this diluted sample was transferred to a 50mL volumetric flask and the sample was further diluted to 50 mL with de-ionized H_2_O. The chemical/metallic composition of the samples was then determined by inductively coupled plasma (ICP) mass spectrometry (Thermo Scientific™ iCAP™ 7600 ICP-OES). Data were analyzed using Student’s T performed manually.

### Molecular methods

DNA was isolated using the MoBio Power Soil DNA isolation kit according to manufacturer’s instructions. Nine DNA isolations and subsequent analyses were performed on each of the nine *S. citrinum* and nine *P. arhizus* individuals for a total of 81 individual extractions from each species. To maximize the likelihood that the fruiting bodies sampled were indeed individual “islands” soils surrounding fruiting bodies were screened for the presence of *S. citrinum* and *P. arhizus* using ITS-RFLP and fruiting bodies were collected for analysis only when areas between fruiting bodies lacking these fungi were detected. In addition, we amplified the nuclear Internal Transcribed Spacer region of each *S. citrinum* and *P. arhizus* individual. We used the ITS3/ITS4 primers in the PCR and sequencing reactions and utilized PCR conditions previously described (White et al. 1990). This was undertaken to ensure that we were analyzing single species of both fungi in our study.

The microbiome community compositions *S. citrinum* and *P. arhizus* were determined via 16S rRNA gene sequence amplicon sequencing (itags). For this analysis, PCR reactions were undertaken using sporocarp extractions using the PHusion High-Fidelity PCR Master Mix (New England Biolabs). Libraries were generated using the TruSeq DNA PCR-Free Prep Kit (Illumina) and quality was assessed using Qubit 2.0 on a Thermo Scientific Fluorometer and Agilent Bioanalyzer 2100 system. The library was sequenced using an IlluminHiSeq2500 platform. 16S itags for the V3/V4 regions were generated by Novogene Corporation. Sequences were assembled using FLASH V1.2.7 (Grayston and Wainwright 1988) and data were quality filtered using QIIME V2 using the default parameters (Bolyen et al. 2019). Chimeras were removed using UCHIME (Edgar et al. 2011). Sequences were analyzed using UPARSE v7.0.1001 (Edgar 2013) and sequences with >97% similarity were clustered as OTUs. Multiple sequence alignments were performed using MUSCLE V3.8.31 (Edgar 2004).

Taxonomic annotation was accomplished using the GreenGene Database version 13_8 (included in the QIIME software package mentioned above), based on the RDP Classifier v2.2, and also by using QIIME-compatible SILVA (Yilmaz et al. 2014) and BLAST (Altschul et al. 1997). Alpha and Beta diversity measures described below were performed also using QIIME.

To rule out external contamination, we took swipes from all lab surfaces and from inside the laminar flow hood and clean box used for sample and reagent preparation and subjected these swipes to PCR amplification of 16S rRNA genes. Further, we ran reaction blanks on all individual reagents in the analysis stream. All reagents were negative, and no swipes from any surface produced sequences that matched any of the 16S rRNA gene sequences obtained from *P. arhizus* individuals.

The communities were analyzed using standard ecological measures. Data are reported on a per fruiting body basis. Good’s Coverage was used to estimate the percent of total species represented in the sampling. Anosim (a non-parametric analysis of similarity) and Analysis of Molecular Variance (AMOVA) were used to detect differences in populations based on molecular markers. Bray Adonis was used to further assess variance between populations. Principal Components Analysis was used as a method of orthological transformation to convert possibly correlated data points into principal components which can be visualized and samples grouped graphically. Taxon richness was assessed using CHAO1 and ACE (both being abundance-based estimator of species richness) (Chao 1984; Chao and Chiu 2016) and whole tree phylogenetic diversity (PD Whole Tree) which is a phylogenetic analog of taxon richness and is expressed as number of tree units found in a sample (Chao and Chiu 2014). Rarefied Shannon and Simpson’s Diversity indices were used as diversity measures which account for species abundance, numbers, evenness and richness. The Wilcoxin’s Signed Rank Test (a non-parametric version of the T-test) was used to determine whether the two communities were derived from the same population. The above analyses were undertaken using QIIME as indicated previously in this section.

## Results

Elemental Analyses (Table 1) demonstrated that both *S. citrinum* and *P. arhizus* contain a rich combination of elements. Despite lacking the visible Si/S inclusions present within *P. arhizus* the fruiting bodies of *S. citrinum* are enriched in K, Si and S, and also Zn relative to those of *P. arhizus*. In contrast, the fruiting bodies of *P. arhizus* are enriched significantly relative to *S. citrinum* in Cd, Ca, Co, Cu, Li, Mo, Ni, K, Va.

**Table 1.**
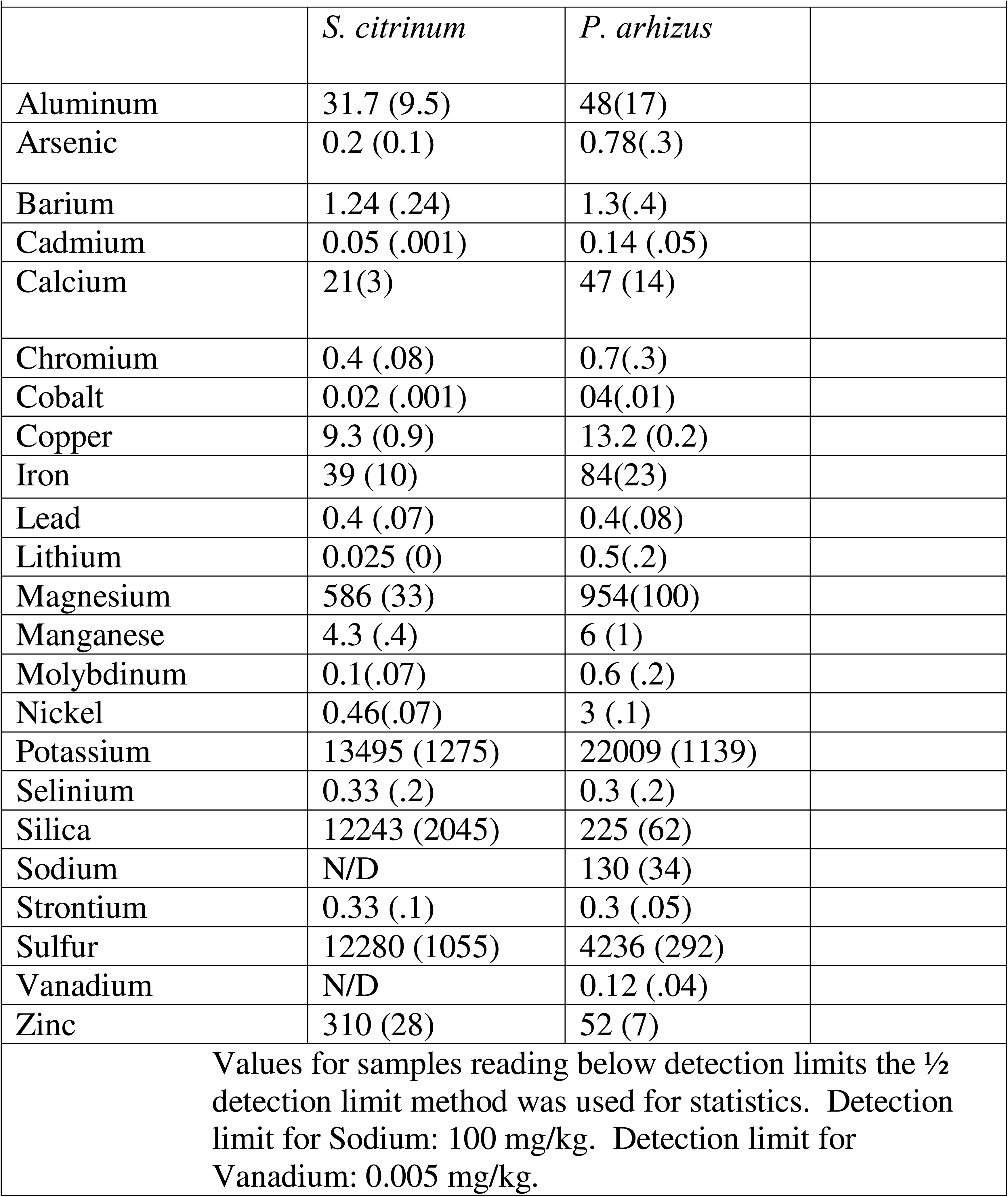
ICP-MSl analysis of S. citrinum and P. arhizus fruiting bodies (n=9, Mean elemental concentration: mg/kg and (SE)

All ITS amplicons from *S. citrinum* and *P. arhizus* specimens sampled in this study differed from each other, and are internally identical (Appendix 1). These data were further supported the uniform observed sporocarp morphologies of the specimens (Figure 1) and are consistent with our previous studies (Cullings and Makhija 2001; Cullings et al. 2020).

Alpha Diversity measures of itag data indicate a high Good’s Coverage (.0.994/0.995 for *S. citrinum* and *P. arhizus* respectively). Statistical analyses demonstrate that the microbiome of *P. ahrizus* is distinct from that of *S. citrinum* (Figure 2), and significantly richer and more diverse as indicated by all alpha diversity measures (Table 2).

**Figure 2:**
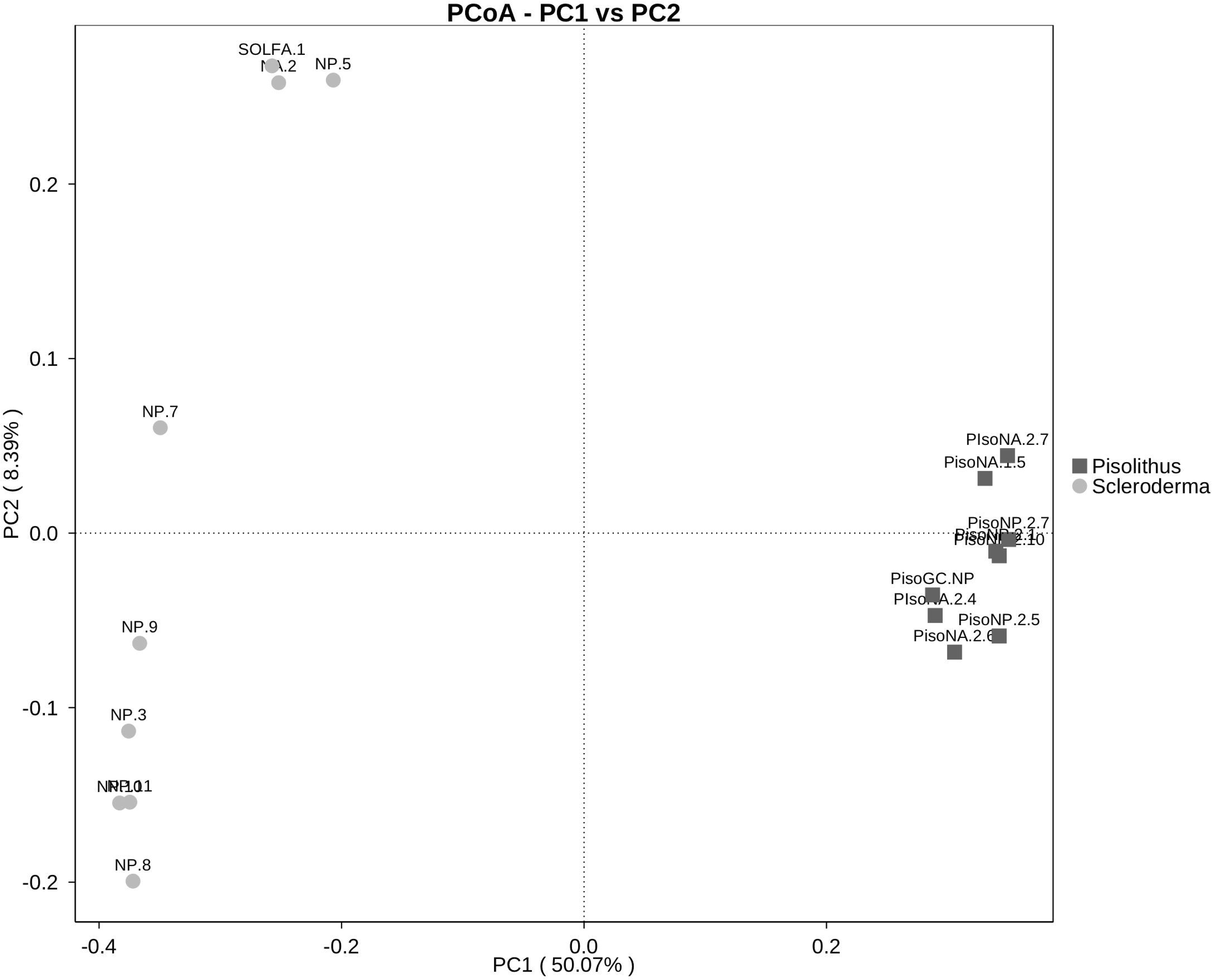
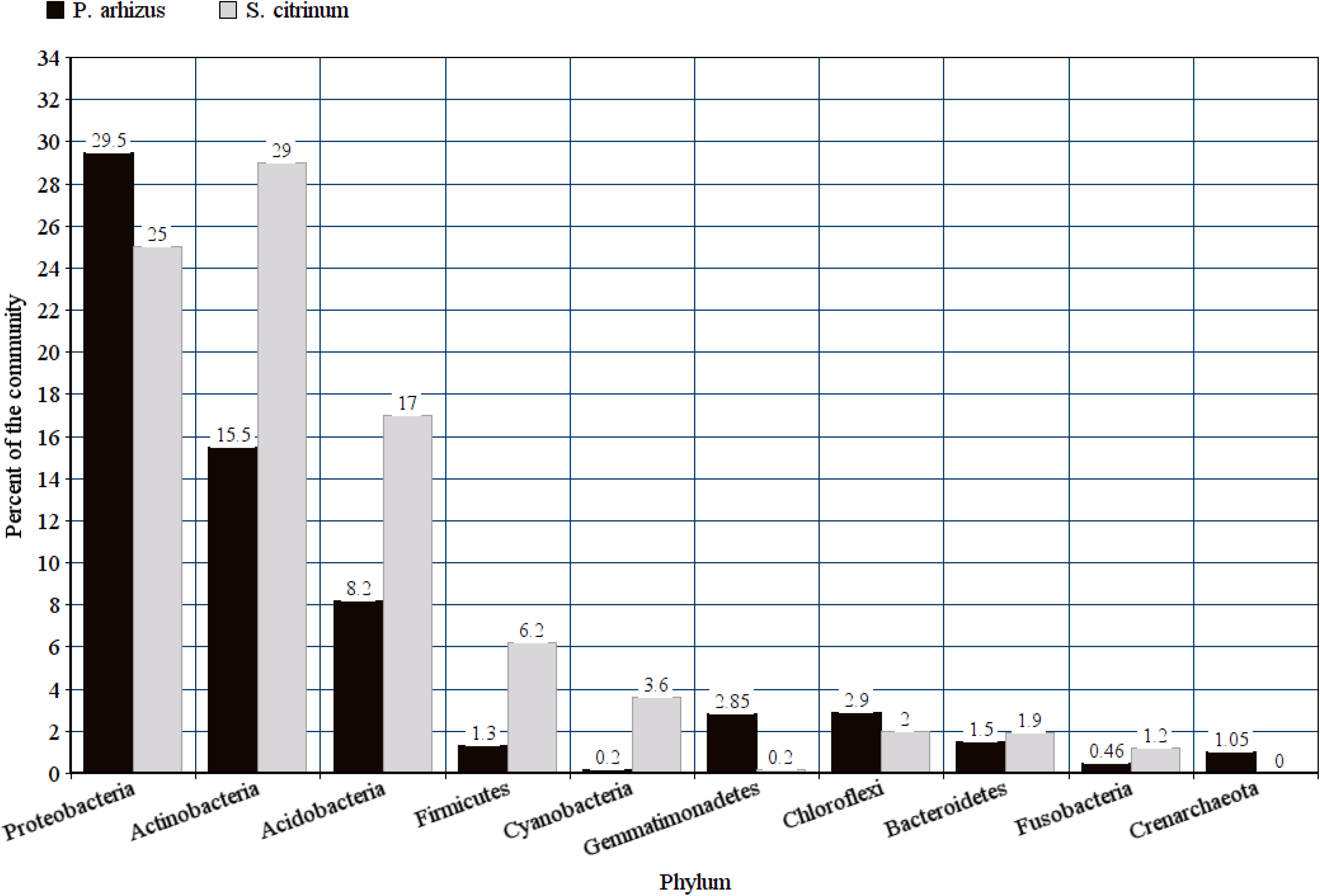
Principal components analysis of the microbiomes of *S. citrina* and *P. arhizus*, indicating that the communities are comprised of different populations of bacteria

**Table 2.** Alpha diversity measures, Mean (SE)

| Table 2. Alpha diversity measures, Mean (SE) |  |  |  |
| --- | --- | --- | --- |
|  | *S. citrinum* | *P. arhizus* | P< |
| Rarefied Shannon Diversity | 6.905 (0.08) | 8.986 (0.25) | 0.0001 |
| Simpson's Diversity | 0.974 (0.003) | 0.990 (0.002) | 0.0006 |
| Chao-1 | 947.02 (34.1) | 2224.1 (24.7) | 0.0001 |
| PD Whole tree | 155.50 (13.1) | 187.68 (6.77) | 0.0001 |
| ACE | 1204.31 (17.6) | 2224.91 (26.83) | 0.0001 |

Taxonomic comparisons at the phylum level demonstrate that the microbiomes of *S. citrinum* and *P. arhizus* are dominated by 10 taxa that comprise at least 1% of the microbiome in either or both species; Proteobacteria, Actinobacteria, Acidobacteria, Gemmatimonadetes, Chloroflexi, Bacteroidetes, Crenarchaeota, Firmicutes, Cyanobacteria (chloroplast, Appendix 1 for sequence), and Fusobacteria, (Figure 3a). Statistical analysis demonstrates that the microbiomes are significantly different with that of *S. citrinum* being enriched in Cyanobacteria, Saccharibacteria (TM-7), and Planctomycetes, while that of *P. arhizus* is enriched in Gemmatimonadetes, Latescibacteria, Elusomicrobia, and Tectomicrobia (Figure 3b).

**Figure 3:**
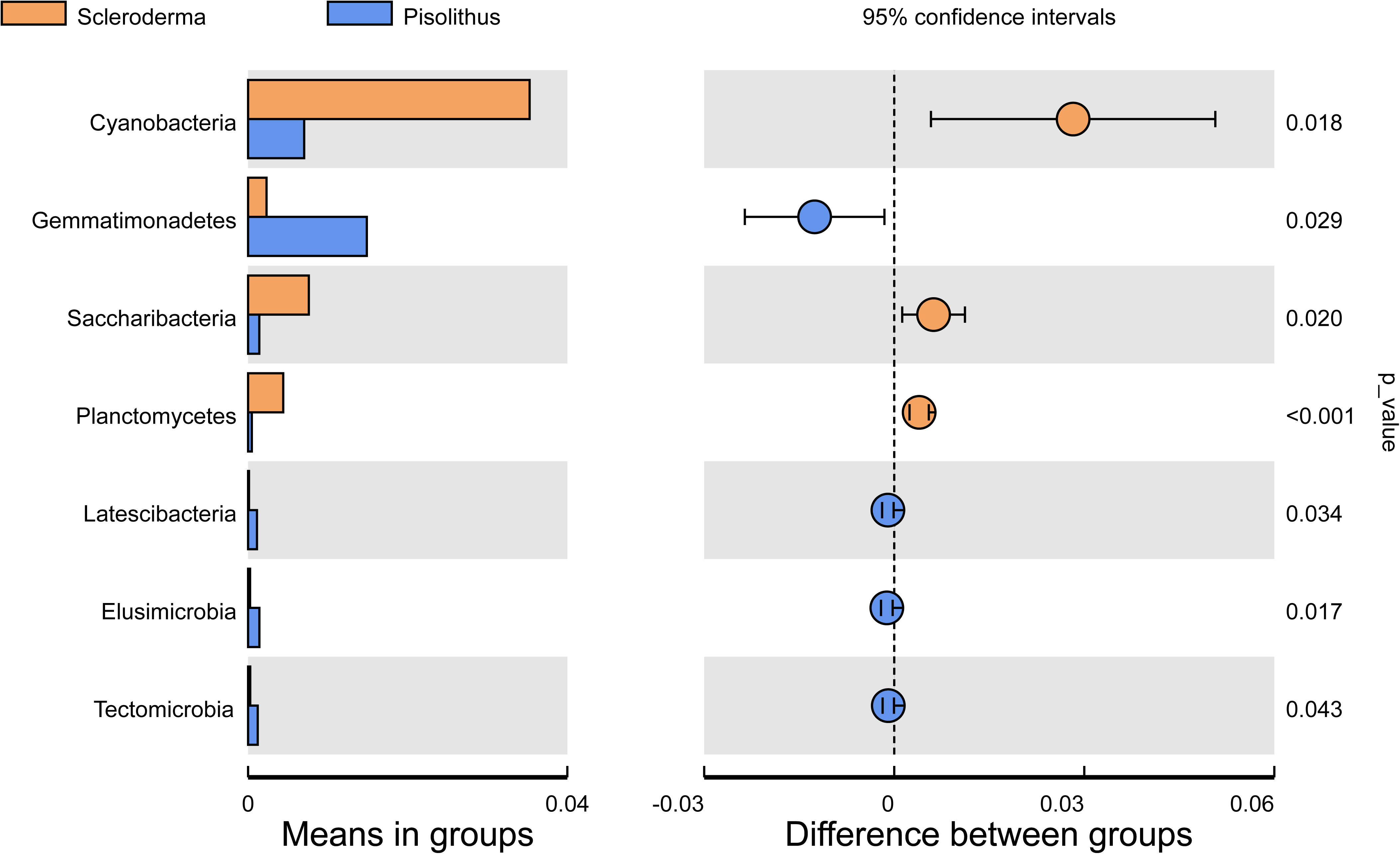
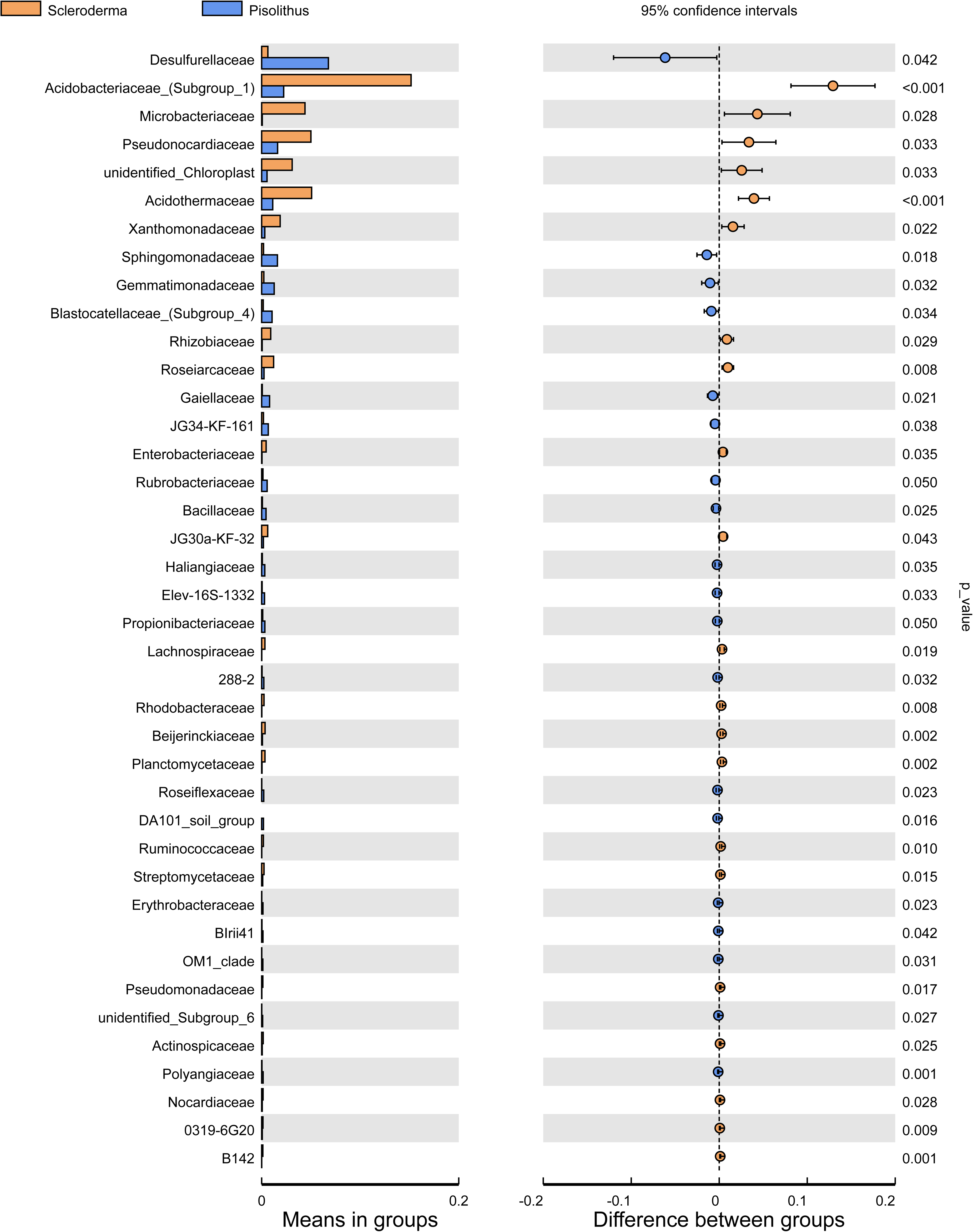
Phylum level abundances and statistical analysis comparison of the *P. arhizus* microbiomes to soil microbial communities. a: comparative relative abundances of microbial phyla inhabiting the microbiomes of *S. citrina* and *P. arhizus*, b: Wilcoxin’s test results of community differences at the phylum level and c, the family level.

At the family level (figure 3c), *S. citrinum* is enriched in Acidobacteriaceae (phylum Acidobacteria driven primarily by the genus *Acidobacterium*, OTU 132, A2, figure A3); Microbacteriaceae (Actinobacteria, driven primarily by the genus *Frigoribacterium* (e.g., OTU 19, A2); Pseudonocardiaceae (Actinobacteria, driven primarily by the genus *Crossiella* (OTU 10, A2); the unidentified chloroplast (Cyanobacteria); the Acidothermaceae (Actinobacteria, driven primarily by the genus *Acidothermus*, OTU 29, A2); the Xanthomonadaceae (Proteobacteria); the Rhizobiaceae (Proteobacteria, driven primarily by genus *Rhizobium* (e.g., OTU 83, A2); and the Roseiarcaceae (Proteobacteria, driven primarily by *Roseiarcus* (e.g., OTU 107, A2).

In contrast, the microbiome of *P. arhizus* is significantly enriched in Desulfurellacea (Proteobacteria, driven primarily by H16, OTU 568, A2); Sphingomonadaceae (Proteobacteria, driven primarily by the genus *Sphingomonas* (e.g., OTU 37, A2); Gemmatimonadaceae (Gemmatimonadetes), Blastocatellaceae (Acidobacteria), Gaiellaceae (Actinobacteria, driven primarily by the genus *Gaiella*, e.g., OTU 96, A2); JG34-KF-161 of the Proteobacteria; the Rubrobacteriaceae (Actinobacteria, driven primarily by the genus *Rubrobacter (*e.g., OTU 54, A2); and the Bacillaceae (Firmicutes). Each housed several additional families that were present in very low frequency, representing less than 0.1% of the overall communities and not discussed individually.

## Discussion

Our data indicate that the microbiomes of both *P. arhizus* and *S. citrinum* are comprised of a diverse assemblage of microbes. However, despite being identical in terms of mode of C/N nutrition (thought to be a primary determinant of endofungal bacterial community composition, Pent et al. 2020) and encountering identical ambient soil conditions (Cullings et al. 2020), their microbiomes differ significantly in richness, diversity and composition.

The microbiome of *P. ahrizus* is both richer and more diverse than that of *S. citrinum*. Why that might be is open to conjecture, however we hypothesize that one factor that could be contributing to this pattern is the difference visible complexity of the mushroom interior; *P. arhizus* contains visible sulfur/silica inclusions, while the interior of *S. citrinum* is more homogeneous and lacking in any inclusions (Cullings et al. 2020, Figure 1). These inclusions and the chemical interfaces they would create could provide energy transitions that provide for greater complexity in the microbial community. To investigate this possibility, we are planning *in situ* hybridization experiments and designing probes to investigate the visualize the distribution of different microbial groups within the fruiting bodies of both species.

Despite the two fungal species being identical in terms of C/N nutrition, our data indicate significant differences in community composition as well. At the phylum level, analysis indicates cyanobacteria signatures, i.e., chloroplast, is abundant in the *S. citrinum* microbome at levels significantly higher than those detected in *P. arhizus* where they are present but in low abundance. Focused BLAST and SILVA identified this entity as a 99% similarity with 100% coverage to the chloroplast of a Rhodophyta species, *Galdieria sulphuraria*. This species is a unicellular, thermoacidophyllic red alga. It belongs to an ancient lineage common to hot, acidic thermal areas, exhibits high resistance to metal toxicity (e.g., Barbier et al. 2005) and it is capable of photosynthesis while in cryptoendolithic growth (e.g., Gross et al. 1998). The presence of these inside the *S. citrinum* microbome provides a plausible source of fixed carbon in the absence of exogenous carbon that would be obtained were this strain ectomycorrhizal.

Analyses indicate that the Saccharibacteria (TM-7) is also significantly more abundant within *S. citrinum*. This group possesses ultra-small cells (Limos et al. 2019) and is present in a wide range of habitats from soils and sediments, to human (e.g., oral and skin) animal (e.g., mammal gut and sponge) microbiomes, to seawater, wastewater, hexavalent chromium contaminated sites, and to toxic sludge indicating a diverse and flexible functionality in the Candidate Phylum.

However, many lineages exhibit habitat or host specificity (reviewed by Ferrari et al. 2015; Zhang et al. 2020). Analysis utilizing SILVA indicates that the dominant OTU presents 95% similarity (100% coverage) to the order Saccharimonadales, family Saccharimonadaceae (Dinnis et al. 2011; Kindaichi et al. 2016), Saccharibacteria Subdivision 1, a group that is prevalent in such diverse habitats as soil, water and wastewater, activated sludge, and nitrifying acidic batch reactors (Hanada et al. 2014). One study suggests that this subdivision is capable of 1,4-dioxane degradation, though the mechanism is as yet unknown (Nam et al. 2016).

Planctomycetes are also more abundant in the *S. citrinum* microbiome. The dominant Planctomycete OTU (548, A2) could only be classified to Order, CPla-3 termite group, and hence may represent a novel family in this group. Wide ranging and diverse, planctomycetes are found in habitats ranging from the Atacama Desert to hot springs (Fuerst 1995; Fuerst and Sagulenko, 2011; Neef et al. 1998) and can dominate microbial communities in geothermal settings (e.g., Badhai et al. 2015; Herbold et al. 2014; Soo et al. 2009). CPLa-3 are plentiful in the gut microbiota of termites (e.g., Hervé et al. 2020; Köhler et al. 2008) and can also occur in acidic terrestrial environments such as mine drainages and (e.g., Ettamimi et al. 2019; Gavrilov et al. 2019).

In contrast, the microbiome of *P. arhizus* is enriched in the phyla Gemmatimonadetes, Latescibacteria (WS3), and Elusimicrobia relative to that of *S. citrinum*. The Gemmatimonadetes are widespread in nature, and were first discovered in a sewage treatment outflow (Zhang et al. 2003). Members of the Gemmatimonadetes have recently been found as components of deep anoxic seafloor sediments and (Fang et al. 2019), and also in soils and near shore sandy sediments (Conte et al. 2018; Huang et al. 2019). Members of the phylum can contain the genetic pathways to fully reduce sulfate (Baker et al. 2015), a function that would be advantageous in the *P. arhizus* microbiome.

The Latescibacteria (WS3, represented by UTO 369, A2 and which could not be classified below the phylum level), was first discovered in anoxic/hydrocarbon-rich lake sediments (Youssef et al. 2015). These have since been found in varied settings including marine, freshwater, terrestrial, bioremediation and symbiotic settings (Markowitz et al. 2014), and members of this group have been detected in metagenomes of anoxygenic microbial mats in YNP. Metagenomics analyses indicate that their functionality includes genes coding for dockerins within cellulosomes where these gene assemblies would enable breakdown of complex hydrocarbons (Farag et al. 2017), again a function that would be advantageous in the rich hydrocarbon environment that we have characterized previously within the fruiting bodies of *P. arhizus* (Cullings et al. 2020).

The Elusimicrobia (OTU 445, classified by SILVA and BLAST only to order, Lineage IV, A2), is a deeply diverging, widespread phylum previously known as Termite Group 1 (Brune 2015; Geissinger et al. 2009; Herleman et al. 2007). Members of this phylogenetically deeply rooted group are fermentative, obligate anaerobes that are also found in anoxic habitats that include contaminated and toxic aquifers. Lineage IV contains novel genes coding for nitrogen fixation (e.g., Méheust et al. 2019) and genes encoding radical S-adenosylmethionine (SAM) 213 proteins which perform many functions including methylation, anaerobic oxidation, and antibiotic biosynthesis (Sofia et al. 2001).

And finally, the Tectomicrobia. The most abundant OTU (350, A2) could not be classified beyond phylum, hence its role here is difficult to ascertain. However, known members of this group can be widely distributed and comprised of carbon-fixing symbionts of sponges which contain anaerobic zones within their bodies. Tectomicrobia produce nearly all the bioactive compounds extracted from those organisms (Jaspars and Challis, 2014; Wilson et al. 2014). Genomic analysis indicates the presence of RuBisCO suggesting carbon fixation, as well as genes coding for sulfate reduction (Liu et al. 2016), and transcriptomic indicates the ability to scavenge ammonia (Feng et al. 2018).

At the family level the micribomes of *S. citrina* fruiting bodies are enriched relative to those of *P. arhizus* by several additional bacterial families; the Microbacteriaceae, the Acidobacteriaceae Subgroup 1, the Pseudonorocardiaceae, the Xanthomonadaceae, the Acidothermaceae, the Rhizobiaceae, and the Roseiarcaceae.

The Microbacteriaceae (driven primarily by the genus *Frigoribacterium*) is a diverse group of comprised of aerobic psychrophiles, soil mesophiles and root endophytes (Dastager et al. 2008; Kämpfer et al 200; Wang et al 2015). The Acidobacteriaceae Subgroup 1 is comprised of aerobic acidophilic, carbon degraders that can be present in the microbiome of metal-rich, uranium-contaminated acidic soils, termite guts, *Sphagnum* peat, acidic mine spoils, as well as metal-rich and (Reviewed by Campbell 2014; Männistö et al. 2012; Falagán et al. 2017).

The Pseudonorocardiaceae (driven primarily by *Crossiella*) is widespread, found in habitats that include terrestrial soil, lake and marine sediments, and the human microbiome (e.g., Zhang et al. 2016) and aerobic thermophiles in geothermal settings (e.g., Ningsih et al. 2019). *Crossiella* is physiologically diverse and has been found in habitats ranging from deep ocean sediments e.g., (Gorzari et al. 2019) to snow packs (e.g., Chuvochina et al. 2011) and karst (e.g., Yun et al. 2016), and in habitats with elevated Cu, Fe, Mn, Zn (e.g., Wisechart et al. 2019). Interestingly, while both *S. citrinum* and *P. arhizus* contain these elements (Table 1), *S. citrinum* is enriched only in Zn relative to *P. arhizus*.

The Xanthomonadaceae is a group that includes radio- and acid-tolerant denitrifyers (e.g., Dahal and Kim 2017; Green et al. 2012; Prakash et al. 2012; Van Den Heuvel et al. 2010). Some strains are capable of mobilizing xenobiotics (e.g., Kanaly et al. 2002), and some can grow in biofilm on the surfaces of sulfur particles (e.g., Lee et al. 2006). The Acidothermacea is a single-genus family (*Acidothermus*) that is comprised of thermophilic, moderately acidophilic, sulfur-oxidizing cellulolytic bacteria of hot springs (Mohagheghi et al. 1986), and is capable of xenobiotic metabolism under extremely harsh conditions e.g., (Devpura et al. 2017; Li et al. 2016).

The Rhizobiaceae, driven primarily by the genus *Rhizobium*, is a soil and nodulating, nitrogen-fixing genus of the Rhizobiaceae that is particularly prone to horizontal gene transfer and can promote plant growth via production of phytohormones gibberellins, auxins and cytokinins. They produce exopolysaccharides (EPS), which can convey resistance to desiccation and antiobiotics (reviewed by Alves et al. 2014). *Rhizobium* strains are being investigated for their industrial uses which include biosorption of heavy metals (particularly Co, Cd), radioactive metals, and biopolymer emulsification.

The Roseiarcaceae, driven primarily by *Roseiarcus*, is a moderately acidophilic, methanotrophic, bacteriochlorophyll a-containing fermentative bacterium capable of chemoautotrophic growth and nitrogen fixation (e.g., Kulichevskay et al. 2014). This group may be a mycorrhizal helper bacterium of another basidiomycete mushroom-forming fungal genus, *Russula* (e.g., Yu et al. 2010) and hence may be predisposed to forming symbioses with basidiomycete fungi.

In contrast, the microbial families that are enriched in the *P. arhizus* fruiting bodies and present in greater than 1% of the total community include the Desulfurallaceae, the Sphingomonadaceae, the Gaiellaceae, the Gemmatimonadaceae the Blastocatellaceae subgroup 4, Alphaproteobacteria family JG34-kf-161, the Bacillaceae, the Rubrobacteriaceae and the Enterobacteriaceae.

Bacteria of the family Desulfurellaceae (driven primarily by H16) are typically strict anaerobic sulfur-reducers, and can also be abundant at sites with high redox potential (> 400 mV) and in the presence of oxygen (> 4 mgL-1) (Reviewed by Greene 2014).

The Sphingomonadaceae (driven primarily by the genus *Sphingomonas*) is a group comprised of sulfur-oxidizers that can be effective in PAH degradation and desulfurization of aromatic hydrocarbons and other xenobiotics. As such, *Sphingomonas* strains are of great interest to the bioremediation community. (e.g., Chen et al. 2008; Gunam et al. 2006; Guo et al. 2010; Li et al. 2005; Li et al. 2007; Zhao and Wang 2012). This set of functions makes sense given the sulfur content and rich hydrocarbon metabolome we have detected within the *P. arhizus* fruiting bodies (Cullings et al. 2020 The Gaiellaceae (driven primarily by *Gaiella*) is a phylogenetically deeply branching, aerobic, nitrate reducing, chemoorganotrophic actinobacterium (Alburquerque et al. 2011; da Costa 2014; Severino et al. 2019), that can be dominant in water-stressed environments (e.g., Sivakala et al. 2017).

The family Gemmatimonadaceae, is comprised of anaerobic thermophiles that can be effective in organic pollutant and petrochemical removal in xenobiotic-contaminated settings (Wagner and Wiegel 2008; Ding et al. 2018).

The Blastocatellaceae subgroup 4, is a group that is comprised of aerobic, chemoorganotrophic, acidophilic mesophiles (e.g., Pascual et al. 2015).

The Alphaproteobacteria family JG34-kf-361 is a group of extremophiles that has been found in habitats such as soil crusts (Liu et al. 2017) and as symbionts of lichen (Aschenbrenner et al. 2017) and as a component of bacterial communities inhabiting uranium mine spoils (Selenska-Pobell et al. 2002).

The Bacillaceae of the Firmicutes is found in a wide array of extreme environments where they can function as radio-tolerant, mercury-resistant anaerobic and aerobic chemolithotrophs, iron-sulfur, Mn-oxidizers. They are present as aerobic thermophiles, found in sulfur hot springs and hydrothermal vents, globally (Logan and Allan 2008, Romano et al. 2005). They can be anaerobic iron- and Mn-reducers in thermal aquifers, mercury-resistant chemolithotrophs in geothermal springs (Chatziefthimiou et al. 2007), sulfur- and hydrogen-oxidizers in thermophilic environments (Beffa et al. 1996).

The Rubrobacteriaceae, driven primarily by the genus *Rubrobacter* (e.g., OTU 54, A2). *Rubrobacter* can be radio- and halotolerant thermophiles, sometimes found in hot springs and of interest for bioremediation of pollution streams (Carreto et al. 1996; Chen et al. 2004; Empadinhas et al. 2007; Ferreira et al. 1999).

And, the Enterobacteriaceae, is a diverse group comprised of facultatively anaerobic nitrate reducers and thermophiles. They can be components of gut microbiota, present as parasites of plants and animals, and denizens of both water and soil habitats. (e.g., Aditiawati et al. 2009; Khiyama et al. 2012; Koskinen et al. 2008).

The data presented here indicate that the fruiting bodies of both *S. citrinum* and *P. ahrizus* provide an above-ground haven for the growth, easy collection, detection and possible culturing of microbes (both aerobic and anaerobic) that because they are either rare or absent from parent soils (Cullings et al. 2020) would otherwise avoid our detection. The data indicate that the microbiome of *P. arhizus* is both richer and more diverse than that of *S. citrinum*, and that the microbiomes of both *S. citrina* and *P. arhizus* are comprised of differential populations photsynthetic, metal-tolerant, aromatic hydrocarbon-degrading microbes that provide plausible sources of energy in the absence of fixed carbon from an ectomycorrhizal host. Further, the microbiome of *P. arhizus* is enriched in anaerobic bacteria relative to those of *S. citrina*, a pattern, which is perhaps not surprising given the known anoxic zones and sharp chemical transitions we have detected and observed within *P. arhizus* fruiting bodies. What processes are driving these diversity and richness patterns is open for speculation. However, our data indicate that while both *S. citrinum* and *P. arhizus* accumulate an array of elements in their fruiting bodies. *S. citrinum* concentrates Si and S to a significantly higher degree than does *P. arhizus*, despite the presence of obvious Si inclusions and elemental sulfur that are present in the fruiting bodies of the latter. We hypothesize that his pattern of elemental composition and distribution, combined with the energy transitions that would be created across the Si/S and aerobic/anaerobic interfaces are likely creating additional complexity within the *P. arhizus* fruiting body, and thereby contributing to the differential patterns of microbial diversity and richness that we measured.

In addition to the metal-tolerant bacteria mentioned above, microbiome communities of both fungal species also contain radio-tolerant microbial strains. Our elemental analyses did not include measures of radioactive isotopes, however *P. tinctorius* (a close relative of *P. arhizus* (Lebel et al. 2018) is known to accumulate uranium (U), thorium (Th) and neodynum (Nd) (Campos et al. 2009). The presence of bacterial families that contain radio-tolerant strains would seem to indicate that the fruiting bodies of *S. citrinum* could be accumulating radioisotopes as well.

The ability of both *Scleroderma* and *Pisolithus* species to tolerate and to sequester heavy metals and radioisotopes has prompted some to put forth both fung*i* as candidates for use in myco- and phyto-remediation strategies targeted at both types of pollutant (e.g., Carrillo-González and González-Chávez 2012; Diagne et al 2015; Kahn et al 2000; Ogo et al. 2017). It is clear that at least some of the metal-remediation abilities are due to functions inherent to the fungus itself (e.g., hexavalent chromium, Shi et al 2018). However, given the scavenging and mineralization abilities of the microbes detected here, one could speculate that the apparent ability of these fungi to serve in this capacity might be due at least as much to the microbial communities housed within as to any inherent abilities on the part of the fungus itself. Targeted co-culture experiments involving the fungus and microbes cultivated from the *S. citrinum* microbiome could help determine if this is the case. At the very least, the easily collectible fruiting bodies of these fungi could provide a rich source of heavy metal- and radiotolerant bacteria for bioremediation use.

Many of the microbes residing within the microbiomes could provide microbial solutions to other ongoing water and soil contamination problems as well. From our previous study of *P. arhizus* we know that the interior of this species contains a metabolome rich in aromatic hydrocarbons and other complex xenobiotic analogs (Cullings et al. 2020). Though in this study we did not investigate the metabolome of *S. citrinum* it is possible, perhaps even probable given the diversity of microbes residing within, that similar compounds also exist within the fruiting bodies of that species. A number of the identified microbial phyla and families are comprised of taxa that include symbionts of termite and ant hind- and mid-gut, and some that apparently thrive on pollution streams and sludge, all of which can be comprised of chemical analogs to the compounds found in the natural substrates utilized by mid- and hindgut-inhabiting microbial symbionts. Because of this, we have begun to address the potential for these fungi to provide bacterial strains for aromatic hydrocarbon remediation use. By using relatively simple methods utilizing fruiting body extracts (adapted from Tillakarathna 2016) we have isolated several novel microbial strains from within *P. arhizus* individuals. Preliminary work in obtaining annotated genomes from these cultures (e.g., Accessions CP066779, CP065858) indicate that they possess the genetic wherewithal to degrade a wide array of petroleum-derived contaminants, and that they contain novel genes that provide enhanced sulfur and metal tolerance. We plan to continue this potentially fruitful line of endeavor.

## Acknowledgements

We would also like to thank Hanna-Annette Peach for sample preparation for the elemental analysis, and for obtaining shipping permits. Data are archived in Genbank under Accessions KEBH00000000, CP066779, CP065858. This work supported by Exobiology grant NASA-Exobiology (13-EXO-007) to Cullings as P.I.

## Appendices

Appendix 1

Consensus ITS-2 reference sequence obtained from *S. citrinum* fruiting-bodies. All ITS-2 sequences obtained in this study were identical.

TTTGAGGTCAACATCGATGACGCGCGCGGCCCGTCCGAGACGGGCTCGCAGAGTTGGAAA GCGACGCGATCCACGCACTTCCAGCCCACGACGGTCATTATGACGTCGAAGAGGCCGTGCCGTGCG AGGCTCGCACACGCCGCTAATGCTTTTGAGGAGAGCCGAAGGTCCCCCTCCCGACAGGAGGGGACC CGCCCGCAGACTCCCATGGTCCAAACCTAGCTCCGACGGGGGTCGAAAGCTTCGATTTGATATTTC GATGACACTCAAACAGGCATGCTCCTCGGAATACCGAGGAGCGCAAGGTGCGTTCAAAGATTCGAT GATTCACGGAAAATCTGCAATTCACATTACTTATCGCGATTCGCTGCGTCCTTCATCGATGCGAGA GCCAAGAGATCCATTGCTGAAAGTTGTACTAGGTTTACGATGGCCGAGGCCACGGACGACATTCTG TAGACATGCGAGTTCTGAGAAGACATAGGTCCCTAAGGACCTACAGCGGGTGCACACGGGTGTGGG AGGTATGAAGCCTCGGAAAGGTTCGAAGGAGCGACGCGAGGACTCTCCCCCTCCCCCTCCCTAGAG GTTCTATTTCGATAATGATCCTTCCGCAGGTTCACCCTACGGAAG

Consensus ITS-2 reference sequence obtained from *P. arhizus* fruiting-bodies. All ITS-2 sequences obtained in this study were identical.

GAATCGCGATAAGTAATGTGAATTGCAGATTTTCCGTGAATCATCGAATCTTTGAACGCA CCTTGCGCTCCTTGGTATTCCGAGGAGCATGCCTGTTTGAGTGTCATCGAAATCTCAAACCAAGCT TTTCTTGACTTCGGTCGAAGGCTCGGGTTTGGACCGTTGGGAGTCTGCGGGCGACGCATGTCGTCG GCTCTCCTGAAATGCATTAGCGGTGGGCATGCAAGTCTTGCTTGGCACCAGCCTCTCCCGGCGTCA TAGTGATCGTCGCGGGCTGTCCAGCTGCAAGGGACATGTCCCATGCTTCTCCAACTTTGCGAGCCC TCTCCTGGGCTCTGCGTTCGAAGGCTTGACCTCAAATCAGGTAGGACTACCCGCTGAACTTAA

Appendix 2: Concensus OTUs of genera driving family-level differences

>OTU_10 TGGGGAATCTTGCGCAATGGGCGAAAGCCTGACGCAGCGACGCCGCGTGGgggATGACGG

CCTTCGGGTTGTAAACCTCT TTCAGCCCTGACGAAGCGAAAGTGACGGTAGGGGCAGAAGAAGCACCGGCCAACTACGTG

CCAGCAGCCGCGGTAATACG TAGGGTGCGAGCGTTGTCCGGAATTATTGGGCGTAAAGAGCTCGTAGGCGGCCTGTCGCG

TCGGCTGTGAAAACTTGGGG CTCAACCCTGAGCTTGCAGTCGATACGGGCAGGCTAGAGTTCGGCAGGGGAGACTGGAAT

TCCTGGTGTAGCGGTGAAAT GCGCAGATATCAGGAGGAACACCGGTGGCGAAGGCGGGTCTCTGGGCCGATACTGACGCT

GAGGAGCGAAAGCGTGGGGA GCGAACAGG

>OTU_19 TGGGGAATATTGCACAATGGGCGAAAGCCTGATGCAGCAACGCCGCGTGAGGGACGACGG

CCTTCGGGTTGTAAACCTCT TTTAGTAGGGAAGAAGCGAAAGTGACGGTACCTGCAGAAaaaGCACCGGCTAACTACGTG

CCAGCAGCCGCGGTAATACG TAGGGTGCAAGCGTTGTCCGGAATTATTGGGCGTAAAGAGCTCGTAGGCGGTTTGTCGCG

TCTGCTGTGAAATCTGGggg CTCAACCcccAGCCTGCAGTGGGTACGGGCAGACTAGAGTGCGGTAGGGGAGATTGGAAT

TCCTGGTGTAGCGGTGGAAT GCGCAGATATCAGGAGGAACACCGATGGCGAAGGCAGATCTCTGGGCCGTAACTGACGCT

GAGGAGCGAAAGCATGGGGA GCGAACAGG

>OTU_29 TGGGGAATATTGCGCAATGGGCGAAAGCCTGACGCAGCGACGCCGCGTGGgggATGAAGG

CCTTCGGGTTGTAAACCCCT TTCAGCAGGGACGAAGCGAAAGTGACGGTACCTGCAGAAGAAGCGCCGGCTAACTACGTG

CCAGCAGCCGCGGTAACACG TAGGGCGCAAGCGTTGTCCGGAATTATTGGGCGTAAAGAGCTCGTAGGCGGTCTGTCACG

TCTGCTGTGAAAACTCGGGG CTTAACCCCGAGCCTGCAGTGGATACGGGCAGACTAGAGGTAGGTAGGGGAGAATGGAAT

TCCCGGTGTAGCGGTGAAAT GCGCAGATATCGGGAGGAACACCGGTGGCGAAGGCGGTTCTCTGGGCCTTACCTGACGCT

GAGGAGCGAAAGCGTGGGGA GCGAACAGG

>OTU_37 TAGGGAATCTTGCCCAATGGGCGCAAGCCTGAGGCAGCAACGCCGCGTGGgggACGAAGG

CTTTCGGGTTGTAAACCCCT TTCAGCAGGGACGAAATTGACGGTACCTGCAGAAGAAGCCCCGGCCAACTACGTGCCAGC

AGCCGCGGTAATACGTAGGg ggCGAGCGTTGTCCGGATTTATTGGGCGTAAAGAGCTCGTAGGCGGCTTGATAAGTCGGG

TGTGAAACCTCCAGGCTCAA CCTGGAGCCGCCATCCGAAACTGTCATGGCTAGAGTCCGGTAGGGGATCACGGAATTCCT

GGTGTAGCGGTGAAATGCGC AGATATCAGGAGGAACACCGATGGCGAAGGCAGTGATCTGGGCCGGTACTGACGCTGAGG

AGCGAAAGCGTGGGGAGCGA ACAGG

>OTU_54 CCAGGAATCTTGGGCAATGGGCGAAAGCCTGACCCAGCAACACCGTGTGGGCGATGAAGG

CCTTCGGGTCGTAAAGCCCT GTTGATAGGGACGAAGGGCGAAGGGTTAATAGCCCCTAGCCTGACGGTACCTTTCGAGGA

AGCCCCGGCTAACTACGTGC CAGCAGCCGCGGTAATACGTAGGgggCGAGCGTTGTCCGGAATTATTGGGCGTAAAGAGC

GTGTAGGCTGTTCGGTAAGT CTGCCGTGAAAACCTGAGGCTCAACCTCGGGCGTGCGGTGGATACTGCCGGGCTAGAGGA

CGGTAGAGGCGAGTGGAATT

CCCGGTGTAGCGGTGAAATGCGCAGATATCGGGAGGAACACCAGTAGCGAAGGCGGCTCG CTGGGCCGTTCCTGACGCTG

AGACGCGAAAGCTAGGGGAGCGAACAGG

>OTU_83 TGGGGAATATTGGACAATGGGCGCAAGCCTGATCCAGCCATGCCGCGTGAGTGATGAAGG

CCCTAGGGTTGTAAAGCTCT TTCACCGGAGAAGATAATGACGGTATCCGGAGAAGAAGCCCCGGCTAACTTCGTGCCAGC

AGCCGCGGTAATACGAAGGg ggCTAGCGTTGTTCGGATTTACTGGGCGTAAAGCGCACGTAGGCGGATCGATCAGTCAGG

GGTGAAATCCCAGGGCTCAA CCCTGGAACTGCCTTTGATACTGTCGATCTGGAGTATGGAAGAGGTAAGTGGAATTCCGA

GTGTAGAGGTGAAATTCGTA GATATTCGGAGGAACACCAGTGGCGAAGGCGGCTTACTGGTCCATTACTGACGCTGAGGT

GCGAAAGCGTGGGGAGCAAA CAGG

>OTU_96 TGGGGAATCTTGCGCTAATGCGGGAAACCGTGACGCAGCGACGCCGCGTGAGGGAAGAAG

GCCTTCGGGTTGTAAACCTC TTTCAGGAGGGACGAAGCTACTCGGGTTAATAGCCCAGAGGGTGACGGTACCTCCAGAAG

AAGCCCCGGCTAACTACGTG CCAGCAGCCGCGGTAATACGTAGGgggCAAGCGTTGTCCGGATTTATTGGGCGTAAAGAG

CGTGTAGGCGGCTTTACAGG TCCGTTGTGAAAACTCGAGGCTCAACCTCGAGACGCCGGTGGAAACCGTAAAGCTAGAGT

CCGGAAGAGGAGAGTGGAAT TCCTGGTGTAGCGGTGAAATGCGCAGATATCAGGAAGAACACCCGTGGCGAAGGCGGCTc tctGGGACGGTACTGACGCT GAGACGCGAAAGCGTGGGGAGCGAACAGG

>OTU_107 TGGGGAATATTGGACAATGGGCGCAAGCCTGATCCAGCCATGCCGCGTGAGTGATGACGG

CCTTAGGGTTGTAAAGCTCT TTCGCTCGGGACGATAATGACGGTACCGAgagaAGAAGCCCCGGCTAACTTCGTGCCAGC

AGCCGCGGTAATACGAAGGg ggCTAGCGTTGTTCGGAATTACTGGGCGTAAAGCGCGTGTAGGCGGGTCTTTAAGTCAGG

GGTGAAATGCCAAGGCTCAA CCTTGGAACTGCCTTTGATACTGGAGATCTTGAGTCCGGGAGAGGTGAGTGGAATTGCGA

GTGTAGAGGTGAAATTCGTA GATATTCGCAAGAACACCAGTGGCGAAGGCGGCTCACTGGCCCGGAACTGACGCTGAGAC

GCGAAAGCGTGGGGAGCAAA CAGG

>OTU_132 TGGGGAATTTTGCGCAATGGgggAAACCCTGACGCAGCAACGCCGCGTGGAGGATGAAGT

CCCTTGGGACGTAAACTCCT TTCGACCGGGACGATAATGACGGTACCGGAAGAAGAAGCCCCGGCTAACTTCGTGCCAGC

AGCCGCGGTAATACGAGGgg ggCGAGCGTTGTTCGGAATTATTGGGCGTAAAGGGTGCGTAGGCGGTTCGGTAAGTTTTG tgtgAAATCTTCGGGCTCAA CTCGAAGCCTGCACAAAATACTGCCGGGCTTGAGTGTGGGAGAGGTGAGTGGAATTTCCG

GTGTAGCGGTGAAATGCGTA GATATCGGAAGGAACACCTGTGGCGAAAGCGGCTCACTGGACCACAACTGACGCTGATGC

ACGAAAGCTAGGGGAGCAAA CAGG

>OTU_350 TGGGGAATCTTGGGCAATGGgggAAACCCTGACCCAGCGACGCCGCGTGGgggATGAAGG

CTTTCGGGTCGTAAACCCCT GTTAGGTGGGACGAATGTCCcccTGCAAATAGCTGGgggATTTGACGGTACCACCAGAGA

AAGCCCCGGCTAACTCCGTG CCAGCAGCCGCGGTAATACGGAGGgggCGAGTGTTGTTCGGAATTACTGGGCGTAAAGGG

CgcgcAGGCGGCCTGGCAAG

TCGAGTGTGAAATCCCTTGGCTTAACTGAGGAATGGCGCTCGAAACTACCTCGCTGGAGG GCGGGAGAGGGAAGTGGAAC

TCTCGGTGTAGGGGTGAAATCTGTAGATATCGAGAGGAACACCGGTGGCGAAGGCGGCTT CCTGGCCCGTACCTGACGCT

GAGGCGCGAAAGCGTGGGGAGCAAACAGG

>OTU_369 TCGAGAGGCTTCGGCAATGGgggAAACCCTGACCGAGCGACGCCGCGTGGAGGATGAAGG

CCCTTGGGTTGTAAACTCCT TTCGTGGgggAAGAATGTATTCGAGGTAAATAATCTCGAGTATTGACGGTACCCCAAGAA

GAAGCTCCGGCTAACTCCGT GCCAGCAGCCGCGGTAATACGGggggAGCAAGCGTTGTTCGGAATCACTGGGCGTAAAGG

GAGTCTAGGCGGTTTGGTAA GTGGGATGTGAAATGCCCGGGCTCACCCCGGGACCTGCATCCCAAACTGCTGAGCTTGAG

TACAGGAGAGGGTGGgggAA TTCCAGGTGTAGCGGTGAAATGCGTAGATATCTGGAGGAACACCGGTGGCGAAGGCGCCC

ACCTGGCCTGCTACTGACGC TGAAGCTCGAAAGCTAGGGGAGCAAACGGG

>OTU_445 TGGGGAATTTTGGACAATGGGCGAAAGCCTGATCCAGCAACGCTGCGTGAGGGATGAAGG

TCTTCGGATCGTAAACCTCT TttttGGgggACGAATACCCGCAAGGGTTTGACGGTACTCCAAGAATAAGCCACGGCTAA

ATATGTGCCAGCAGCCGCGG TAAGACATATGTGGCGAGCGTTACTCGGAATTACTAGGCGTAAAGCGCGTGTAGGCGGAT

GCTTAAGTCTGCTGTGAAAT CTCCCGGCTTAACTGGGAGCGGTCAGCGGATACTGGGCGTCTTGAGTGAGGTAGGgggTG

CTGGAATTCCCGGTGTAGCG GTGAAATGCGTAGATATCGGGAGGAACACCGATGGCGAAAGCAGGCACCTGGGCCTTTAC

TGACGCTGAGACGCGAAAGC TAGGGGAGCAAACAGG

>OTU_548 CAACGAATCTTCCGCAATGGGCGCAAGCCTGACGGAGCGACGCCGCGTGTGGGTTAAGTT

CTTCGGAATGTAAACCACTG TTAGGGTTATGAAAGCGATGGAGGATAATATACTCCAAAGTTGATCTAGCCCAGAGAAAG

GGACGGCTAACTCTGTGCCA GCAGCCGCGGTAATACAGAGGTCCCAAGCGTTACTGAGATTCACTGGGTTTAAAGGGTGC

GTAGGTGGTGCGTTAAGTCA GTTGTGAAATCCCCGGGCTCAACCCGGGAATAGCTTCTGATACTGGCGTGCTTGAGGCCG

GTATAGGTTACTGGAACGGA CGGTGGAGCGGTGAAATGCGTAGATATCGTCTGGAACGCCAGTGGTGAAGACGGGTAACT

AGGCCGGTTCTGACACTGAG GCACGAAAGCGTGGGGAGCGAACGGG

>OTU_568 TGGgggATATTGCGCAATGGGCGAAAGCCTGACGCAGCGACGCCGCGTGGGTGATGAAGG

CCTTCGGGTCGTAAAGCCCT GTCGAGGGGAAAGAAACGGTCTCGACCTAACACGTCGGGGCCCTGACGGTACCCCTAAAG

GAAGCACCGGCTAACTCCGT GCCAGCAGCCGCGGTAATACGGAGGGTGCGAGCGTTGTTCGGAATCACTGGGCGTAAAGG

GCGTGTAGGCGGTCCGCTAA GTCTGATGTGAAAGCCCGGGGCTCACCcccGGAAGGGCATTGGAAACTGACGGACTCGAG

TCCCGAAGAGGAGGGTGGAA TTCCTGGTGTAGCGGTGAAATGCGTAGATATCAGGAGGAACACCGGTGGCGAAGGCGGCC

CTCTGGACGGTGACTGACGC TGAGACGCGAAAGCGTGGGGAGCAAACAGG

Appendix 3: Wilcoxin’s test comparisons of the microbiomes of *S. citrinum* and *P. arhizus*.0.2

## References

Aditiawati, P., Yohandini, H., and Fida Madayanti, A. 2009. Microbial diversity of acidic hot spring (Kawah Hujan B) in geothermal field of Kamojang area, west Java-Indonesia. The Open MicrobiologyJjournal 3: 58–66.

Aggangan, N., Pampolina, N., Cadiz, N., and Raymundo, A. 2015. Assessment of plant diversity and associated mycorrhizal fungi in the mined-out sites of Atlas mines in Toledo city, Cebu for bioremediation. Journal of Environmental Science and Management 18(1): 71–86.

Albuquerque, L., França, L., Rainey, F. A., Schumann, P., Nobre, M. F., and da Costa, M. S. 2011. Gaiella occulta gen. nov., sp. nov., a novel representative of a deep branching phylogenetic lineage within the class Actinobacteria and proposal of Gaiellaceae fam. nov. and Gaiellales ord. nov. Systematic and Applied Microbiology 34(8): 595–599.

Altschul, S.F., Madden, T.L., Schäffer, A.A., Zhang, J., Zhang, Z., Miller, W. and Lipman, D.J. 1997. Gapped BLAST and PSI-BLAST: a new generation of protein database search programs. Nucleic Acids Research 25(17): 3389–3402.

Alves, L.M.C., De Souza, J.A.M., de Mello Varani, A. and de Macedo Lemos, E.G. 2014. The family rhizobiaceae. The Prokaryotes, pp.419–437.

Aschenbrenner, I. A., Cernava, T., Erlacher, A., Berg, G., and Grube, M. 2017. Differential sharing and distinct co-occurrence networks among spatially close bacterial microbiota of bark, mosses and lichens. Molecular ecology 26(10): 2826–2838.

Badhai, J., Ghosh, T. S., and Das, S. K. 2015. Taxonomic and functional characteristics of microbial communities and their correlation with physicochemical properties of four geothermal springs in Odisha, India. Frontiers in Microbiology 6: 1166.

Bahram, M., Vanderpool, D., Pent, M., Hiltunen, M., and Ryberg, M. 2018. The genome and microbiome of a dikaryotic fungus (*Inocybe terrigena*, Inocybaceae) revealed by metagenomics. Environmental Microbiology Reports 10(2): 155–166. doi:10.1111/1758-2229.12612

Baker, B. J., Lazar, C. S., Teske, A. P., and Dick, G. J. 2015. Genomic resolution of linkages in carbon, nitrogen, and sulfur cycling among widespread estuary sediment bacteria. Microbiome 3(1): 14. doi:10.1186/s40168-015-0077-6

Barbier, G., Oesterhelt, C., Larson, M.D., Halgren, R.G., Wilkerson, C., Garavito, R.M., Benning, C. and Weber, A.P., 2005. Comparative genomics of two closely related unicellular thermo-acidophilic red algae, Galdieria sulphuraria and Cyanidioschyzon merolae, reveals the molecular basis of the metabolic flexibility of Galdieria sulphuraria and significant differences in carbohydrate metabolism of both algae. Plant Physiology 137(2): 460–474.

Barros, L., Baptista, P., Correia, D. M., Casal, S., Oliveira, B., and Ferreira, I. C. 2007. Fatty acid and sugar compositions, and nutritional value of five wild edible mushrooms from Northeast Portugal. Food Chemistry 105(1): 140–145.

Barros, L., Venturini, B. A., Baptista, P., Estevinho, L. M., and Ferreira, I. C. 2008. Chemical composition and biological properties of portuguese wild mushrooms: a comprehensive study. Journal of Agricultural Food Chemistry 56(10): 3856–3862. doi:10.1021/jf8003114

Beffa, T., Blanc, M., and Aragno, M. 1996. Obligately and facultatively autotrophic, sulfur-and hydrogen-oxidizing thermophilic bacteria isolated from hot composts. Archives of Microbiology 165(1): 34–40.

Benucci, G. M. N., and Bonito, G. M. 2016. The truffle microbiome: Species and geography effects on bacteria associated with fruiting bodies of *Hypogeous pezizales*. Microbial Ecology 72(1): 4–8. doi:10.1007/s00248-016-0755-3

Blaser, M.J., Cardon, Z.G., Cho, M.K., Dangl, J.L., Donohue, T.J., Green, J.L., Knight, R., Maxon, M.E., Northen, T.R., Pollard, K.S. and Brodie, E.L. 2016. Toward a predictive understanding of Earth’s microbiomes to address 21st century challenges. mBio 7(3):00714–16; DOI: 10.1128/mBio.00714-16

Bolyen, E., Rideout, J.R., Dillon, M.R., Bokulich, N.A., Abnet, C.C., Al-Ghalith, G.A., Alexander, H., Alm, E.J., Arumugam, M., Asnicar, F. and Bai, Y. 2019. Reproducible, interactive, scalable and extensible microbiome data science using QIIME 2. Nature biotechnology 37(8): 852–857.

Breznak JA. 1982. Intestinal microbiota of termites and other xylophagousinsects. Annu Rev Microbiol 36:323–343.

Brune, A. 2015. Elusimicrobia. Bergey’s Manual of Systematics of Archaea and Bacteria, 1-3.

Campbell, B. J. 2014. The Family Acidobacteriaceae. The Prokaryotes 405, 415.

Campos, J. A., Tejera, N. A., and Sánchez, C. J. 2009. Substrate role in the accumulation of heavy metals in sporocarps of wild fungi. Biometals 22(5): 835–841.

Carreto, L., Moore, E., Nobre, M. F., Wait, R., Riley, P. W., Sharp, R. J., and Da Costa, M. S. 1996. *Rubrobacter xylanophilus* sp. nov., a new thermophilic species isolated from a thermally polluted effluent. International Journal of Systematic and Evolutionary Microbiology 46(2): 460–465.

Carrillo-González, R. and González-Chávez, M.D.C.A. 2012. Tolerance to and accumulation of cadmium by the mycelium of the fungi *Scleroderma citrinum* and *Pisolithus tinctorius*. Biological trace element research 146(3), pp.388–395.

Cavanaugh, C.M., Gardiner, S.L., Jones, M.L., Jannasch, H.W. and Waterbury, J.B. 1981. Prokaryotic cells in the hydrothermal vent tube worm *Riftia pachyptila* Jones: possible chemoautotrophic symbionts. Science pp.340–342.

Chao, A., 1984. Nonparametric estimation of the number of classes in a population. Scandinavian Journal of Statistics pp.265–270.

Chao, A., and Chiu, C. H. (2014). Species richness: estimation and comparison. Wiley StatsRef: Statistics Reference Online, 1-26.

Cho, Y. S., Kim, J. S., Crowley, D. E., and Cho, B. G. 2003. Growth promotion of the edible fungus *Pleurotus ostreatus* by fluorescent pseudomonads. FEMS Microbiology Letters 218(2): 271–276. doi:10.1016/S0378-1097(02)01144-8.

Chatziefthimiou, A. D., Crespo-Medina, M., Wang, Y., Vetriani, C., and Barkay, T. 2007. The isolation and initial characterization of mercury resistant chemolithotrophic thermophilic bacteria from mercury rich geothermal springs. Extremophiles 11(3): 469–479.

Chen, M.Y., Wu, S.H., Lin, G.H., Lu, C.P., Lin, Y.T., Chang, W.C. and Tsay, S.S., 2004. Rubrobacter taiwanensis sp. nov., a novel thermophilic, radiation-resistant species isolated from hot springs. International Journal of Systematic and Evolutionary Microbiology 54(5): 1849–1855.

Chen, J., Wong, M.H., Wong, Y.S. and Tam, N.F., 2008. Multi-factors on biodegradation kinetics of polycyclic aromatic hydrocarbons (PAHs) by *Sphingomonas* sp. a bacterial strain isolated from mangrove sediment. Marine Pollution Bulletin 57(6-12): 695–702.

Chuvochina, M.S., Alekhina, I.A., Normand, P., Petit, J.R. and Bulat, S.A. 2011. Three events of Saharan dust deposition on the Mont Blanc glacier associated with different snow-colonizing bacterial phylotypes. Microbiology 80(1): 125–131.

Conte, A., Papale, M., Amalfitano, S., Mikkonen, A., Rizzo, C., De Domenico, E., Michaud, L. and Giudice, A.L. 2018. Bacterial community structure along the subtidal sandy sediment belt of a high Arctic fjord (Kongsfjorden, Svalbard Islands). Science of the Total Environment 619: 203–211.

Ceja-Navarro JA, Vega FE, Karaoz U, Hao Z, Jenkins S, Lim HC, Kosina P, Infante F, Northen TR, Brodie EL. 2015. Gut microbiota mediate caffeine detoxification in the primary insect pest of coffee. Nat Commun 6:7618. http://dx.doi.org/10.1038/ncomms8618

Cullings, Ken. and Makhija, S., 2001. Ectomycorrhizal fungal associates of *Pinus contorta* in soils associated with a hot spring in Norris Geyser Basin, Yellowstone National Park, Wyoming. Appl. Environ. Microbiol. 67(12): 5538–5543.

Cullings, K., Stott, M. B., Marinkovich, N., DeSimone, J., & Bhardwaj, S. 2020. Phylum-level diversity of the microbiome of the extremophilic basidiomycete fungus *Pisolithus arhizus* (Scop.) Rauschert: An island of biodiversity in a thermal soil desert. MicrobiologyOpen e1062.

Dahal, R. H., and Kim, J. 2017. Rhodanobacter humi sp. nov., an acid-tolerant and alkalitolerant gammaproteobacterium isolated from forest soil. International Journal of Systematic and Evolutionary Microbiology 67(5): 1185–1190.

Dastager, S. G., Lee, J. C., Ju, Y. J., Park, D. J., and Kim, C. J. 2008. Frigoribacterium mesophilum sp. nov., a mesophilic actinobacterium isolated from Bigeum Island, Korea. International Journal of Systematic and Evolutionary Microbiology 58(8): 1869–1872.

da Costa, L. A. M. S. 2014. 19 The Family Gaiellaceae.

Deveau, A., Bonito, G., Uehling, J., Paoletti, M., Becker, M., Bindschedler, S., Hacquard, S., Hervé, V., Labbé, J., Lastovetsky, O.A. and Mieszkin, S. 2018. Bacterial–fungal interactions: ecology, mechanisms and challenges. FEMS Microbiology Reviews 42(3): 335–352

Devpura, N., Jain, K., Patel, A., Joshi, C. G., and Madamwar, D. 2017. Metabolic potential and taxonomic assessment of bacterial community of an environment to chronic industrial discharge. International Biodeterioration and Biodegradation 123: 216-227.

Diagne, N., Ngom, M., Djighaly, P.I., Ngom, D., Ndour, B., Cissokho, M., Faye, M.N., Sarr, A., Sy, M.O., Laplaze, L. and Champion, A. 2015. Remediation of heavy metal-contaminated soils and enhancement of their fertility with actinorhizal plants. In Heavy Metal Contamination of Soils (pp. 355–366). Springer, Cham.

Ding, P., Chu, L., and Wang, J. 2018. Advanced treatment of petrochemical wastewater by combined ozonation and biological aerated filter. Environmental Science and Pollution Research 25(10): 9673–9682.

Dinis, J.M., Barton, D.E., Ghadiri, J., Surendar, D., Reddy, K., Velasquez, F., Chaffee, C.L., Lee, M.C.W., Gavrilova, H., Ozuna, H. and Smits, S.A. 2011. In search of an uncultured human-associated TM7 bacterium in the environment. PloS one 6(6).

Dubilier, N., McFall-Ngai, M., and Zhao, L. 2015. Microbiology: create a global microbiome effort. Nature 526(7575): 631–634.

Edgar, R. C. 2004. MUSCLE: a multiple sequence alignment method with reduced time and space complexity. BMC Bioinformatics 5: 113. doi:10.1186/1471-2105-5-113

Edgar, R. C., Haas, B. J., Clemente, J. C., Quince, C., and Knight, R. 2011. UCHIME improves sensitivity and speed of chimera detection. Bioinformatics 27(16): 2194–2200. doi:10.1093/bioinformatics/btr381

Edgar, R. C. 2013. UPARSE: highly accurate OTU sequences from microbial amplicon reads. Nature Methods 10(10): 996–998. doi:10.1038/nmeth.2604

Empadinhas, N., Mendes, V., Simoes, C., Santos, M.S., Mingote, A., Lamosa, P., Santos, H. and Da Costa, M.S. 2007. Organic solutes in Rubrobacter xylanophilus: the first example of di-myo-inositol-phosphate in a thermophile. Extremophiles 11(5):.667–673.

Ettamimi, S., Carlier, J. D., Cox, C. J., Elamine, Y., Hammani, K., Ghazal, H., and Costa, M. C. 2019. A meta-taxonomic investigation of the prokaryotic diversity of water bodies impacted by acid mine drainage from the São Domingos mine in southern Portugal. Extremophiles 23(6): 821–834.

Falagán, C., Foesel, B., and Johnson, B. 2017. Acidicapsa ferrireducens sp. nov., Acidicapsa acidiphila sp. nov., and Granulicella acidiphila sp. nov.: novel acidobacteria isolated from metal-rich acidic waters. Extremophiles 21(3): 459–469.

Fang, X. M., Zhang, T., Li, J., Wang, N. F., Wang, Z., and Yu, L. Y. 2019. Bacterial community pattern along the sediment seafloor of the Arctic fjorden (Kongsfjorden, Svalbard). Antonie Van Leeuwenhoek 112(8): 1121–1136. doi:10.1007/s10482-019-01245-z

Farag, I. F., Youssef, N. H., and Elshahed, M. S. 2017. Global distribution patterns and pangenomic diversity of the candidate phylum “Latescibacteria” (WS3). Applied and Environmental Microbiology 83(10). doi:10.1128/AEM.00521-17

Feng, G., Sun, W., Zhang, F., Orlić, S., and Li, Z. 2018. Functional transcripts indicate phylogenetically diverse active ammonia-scavenging microbiota in sympatric sponges. Marine Biotechnology 20(2): 131–143.

Ferrari, B., Winsley, T., Ji, M., and Neilan, B. 2014. Insights into the distribution and abundance of the ubiquitous candidatus Saccharibacteria phylum following tag pyrosequencing. Scientific Reports 4(1): 1–9.

Ferreira, A. C., Nobre, M. F., Moore, E., Rainey, F. A., Battista, J. R., and da Costa, M. S. 1999. Characterization and radiation resistance of new isolates of *Rubrobacter radiotolerans* and *Rubrobacter xylanophilus*. Extremophiles 3(4): 235–238.

Fuerst, J. A. 1995. The planctomycetes: emerging models for microbial ecology, evolution and cell biology. Microbiology 141(7): 1493–1506.

Fuerst, J. A., and Sagulenko, E. 2011. Beyond the bacterium: planctomycetes challenge our concepts of microbial structure and function. Nature Reviews Microbiology 9(6): 403–413.

Fuerst, J. 2004. Planctomycetes: a phylum of emerging interest for microbial evolution and ecology. World Federation for Culture Collections Newsletter 38: 1–11.

Gavrilov, S.N., Korzhenkov, A.A., Kublanov, I.V., Bargiela, R., Zamana, L., Popova, A., Toshchakov, S., Golyshin, P. and Golyshina, O.V. 2019. Microbial communities of polymetallic deposits’ acidic ecosystems of continental climatic zone with high temperature contrasts. Frontiers in Microbiology 10: 1573.

Geib SM, Filley TR, Hatcher PG, Hoover K, Carlson JE, Jimenez-Gasco Mdel M, Nakagawa-Izumi A, Sleighter RL, Tien M. 2008. Lignin degradation in wood-feeding insects. Proc Natl Acad Sci U S A 105: 12932–12937

Geissinger, O., Herlemann, D. P., Mörschel, E., Maier, U. G., and Brune, A. 2009. The ultramicrobacterium “Elusimicrobium minutum” gen. nov., sp. nov., the first cultivated representative of the termite group 1 phylum. Appl. Environ. Microbiol. 75(9): 2831-2840.

Gozari, M., Bahador, N., Jassbi, A. R., Mortazavi, M. S., Hamzehei, S., and Eftekhar, E. 2019. Isolation, distribution and evaluation of cytotoxic and antioxidant activity of cultivable actinobacteria from the Oman Sea sediments. Acta Oceanologica Sinica 38(12): 84–90.

Grayston, S. J., & Wainwright, M. 1988). Sulphur oxidation by soil fungi including some species of mycorrhizae and wood-rotting basidiomycetes. FEMS Microbiology Ecology 4(1): 1–8.

Green, S.J., Prakash, O., Jasrotia, P., Overholt, W.A., Cardenas, E., Hubbard, D., Tiedje, J.M., Watson, D.B., Schadt, C.W., Brooks, S.C. and Kostka, J.E, 2012. Denitrifying bacteria from the genus Rhodanobacter dominate bacterial communities in the highly contaminated subsurface of a nuclear legacy waste site. Appl. Environ. Microbiol. 78(4): 1039–1047.

Greene A.C. 2014. The Family *Desulfurellaceae*. In: Rosenberg E., DeLong E.F., Lory S., Stackebrandt E., Thompson F. (eds) The Prokaryotes. Springer, Berlin, Heidelberg

Gross, W., Küver, J., Tischendorf, G., Bouchaala, N., and Büsch, W. 1998. Cryptoendolithic growth of the red alga *Galdieria sulphuraria* in volcanic areas. European Journal of Phycology 33(1): 25-31.

Gunam, I. B. W., Yaku, Y., Hirano, M., Yamamura, K., Tomita, F., Sone, T., and Asano, K. 2006. Biodesulfurization of alkylated forms of dibenzothiophene and benzothiophene by *Sphingomonas subarctica* T7b. Journal of Bioscience and Bioengineering 101(4): 322–327.

Guo, C., Dang, Z., Wong, Y., and Tam, N. F. 2010. Biodegradation ability and dioxgenase genes of PAH-degrading *Sphingomonas* and Mycobacterium strains isolated from mangrove sediments. International Biodeterioration and Biodegradation 64(6): 419–426.

Hanada, A., Kurogi, T., Giang, N. M., Yamada, T., Kamimoto, Y., Kiso, Y., and Hiraishi, A. 2014. Bacteria of the candidate phylum TM7 are prevalent in acidophilic nitrifying sequencing-batch reactors. Microbes and Environments 29(4): 353–362.

Herbold, C. W., Lee, C. K., McDonald, I. R., and Cary, S. C. 2014. Evidence of global-scale aeolian dispersal and endemism in isolated geothermal microbial communities of Antarctica. Nature Communications 5(1): 1–10.

Herlemann, D. P., Geissinger, O., and Brune, A. 2007. The termite group I phylum is highly diverse and widespread in the environment. Appl. Environ. Microbiol. 73(20): 6682–6685.

Hervé, V., Liu, P., Dietrich, C., Sillam-Dussès, D., Stiblik, P., Šobotník, J., and Brune, A. 2020. Phylogenomic analysis of 589 metagenome-assembled genomes encompassing all major prokaryotic lineages from the gut of higher termites. PeerJ 8: e8614.

Huang, L., Hu, W., Tao, J., Liu, Y., Kong, Z., and Wu, L. 2019. Soil bacterial community structure and extracellular enzyme activities under different land use types in a long-term reclaimed wetland. Journal of Soils and Sediments 19(5): 2543–2557.

Jaspers, M., and Challis, G. 2014. Microbiology: A talented genus. Nature 506(7486): 38–39.

Kindaichi, T., Yamaoka, S., Uehara, R., Ozaki, N., Ohashi, A., Albertsen, M., Nielsen, P.H. and Nielsen, J.L. 2016. Phylogenetic diversity and ecophysiology of Candidate phylum Saccharibacteria in activated sludge. FEMS Microbiology Ecology 92(6).

Jeffries, P. 1999. “Scleroderma. Ectomycorrhizal Fungi Key Genera in Profile.” 187-200.

Jones, M.D. and Hutchinson, T.C. 1988. Nickel toxicity in mycorrhizal birch seedlings infected with *Lactarius rufus* or *Scleroderma flavidum* Uptake of nickel, calcium, magnesium, phosphorus and iron. New Phytol 108: 461–470.

Kämpfer, P., Rainey, F.A., Andersson, M.A., Lassila, E.N., Ulrych, U., Busse, H.J., Weiss, N., Mikkola, R. and Salkinoja-Salonen, M. 2000. *Frigoribacterium faeni* gen. nov., sp. nov., a novel psychrophilic genus of the family Microbacteriaceae. International Journal of Systematic and Evolutionary Microbiology 50(1): 355–363.

Kanaly, R. A., Harayama, S., and Watanabe, K. 2002. Rhodanobacter sp. strain BPC1 in a benzo [a] pyrene-mineralizing bacterial consortium. Appl. Environ. Microbiol. 68(12): 5826–5833.

Kanso, S., Greene, A. C., and Patel, B. K. 2002. Bacillus subterraneus sp. nov., an iron-and manganese-reducing bacterium from a deep subsurface Australian thermal aquifer. International Journal of Systematic and Evolutionary Microbiology 52(3): 869–874.

Khan, A.G., Kuek, C., Chaudhry, T.M., Khoo, C.S. and Hayes, W.J. 2000. Role of plants, mycorrhizae and phytochelators in heavy metal contaminated land remediation. Chemosphere 41(1-2):197–207.

Khiyami, M. A., Serour, E. A., Shehata, M. M., and Bahklia, A. H. 2012. Thermo-aerobic bacteria from geothermal springs in Saudi Arabia. African Journal of Biotechnology 11(17): 4053–4062.

Köhler, T., Stingl, U., Meuser, K., and Brune, A. 2008. Novel lineages of Planctomycetes densely colonize the alkaline gut of soil-feeding termites (*Cubitermes* spp.). Environmental Microbiology 10(5): 1260–1270.

Koskinen, P. E., Beck, S. R., Örlygsson, J., and Puhakka, J. A. 2008. Ethanol and hydrogen production by two thermophilic, anaerobic bacteria isolated from Icelandic geothermal areas. Biotechnology and Bioengineering 101(4): 679–690.

Kulichevskaya, I. S., Danilova, O. V., Tereshina, V. M., Kevbrin, V. V., and Dedysh, S. N. 2014. Descriptions of *Roseiarcus fermentans* gen. nov., sp. nov., a bacteriochlorophyll a-containing fermentative bacterium related phylogenetically to alphaproteobacterial methanotrophs, and of the family Roseiarcaceae fam. nov. International Journal of Systematic and Evolutionary Microbiology 64(8): 2558–256.

Lackner, G., Partida-Martinez, L. P., and Hertweck, C. 2009. Endofungal bacteria as producers of mycotoxins. Trends in Microbiology 17(12): 570–576.

Lebel, T., Pennycook, S., and Barrett, M. 2018. Two new species of *Pisolithus* (Sclerodermataceae) from Australasia, and an assessment of the confused nomenclature of *P. tinctorius*. Phytotaxa 348(3): 163–186.

Lee, C. S., Kim, K. K., Aslam, Z., and Lee, S. T. 2007. Rhodanobacter thiooxydans sp. nov., isolated from a biofilm on sulfur particles used in an autotrophic denitrification process. International Journal of Systematic and Evolutionary Microbiology 57(8): 1775–1779.

Lemos, L. N., Medeiros, J. D., Dini-Andreote, F., Fernandes, G. R., Varani, A. M., Oliveira, G., and Pylro, V. S. 2019. Genomic signatures and co-occurrence patterns of the ultra-small Saccharimonadia (phylum CPR/Patescibacteria) suggest a symbiotic lifestyle. Molecular Ecology 28(18): 4259–4271.

Li, X., He, J., and Li, S. 2007. Isolation of a chlorpyrifos-degrading bacterium, Sphingomonas sp. strain Dsp-2, and cloning of the mpd gene. Research in Microbiology 158(2): 143-149.

Li, Y., Zhao, S., and Wang, Y. 2012. Microbial desulfurization of ground tire rubber by Sphingomonas sp.: a novel technology for crumb rubber composites. Journal of Polymers and the Environment 20(2): 372–380.

Li, D., Sharp, J. O., and Drewes, J. E. 2016. Influence of wastewater discharge on the metabolic potential of the microbial community in river sediments. Microbial Ecology 71(1): 78–86.

Liu, F., Li, J., Feng, G., and Li, Z. 2016. New genomic insights into “Entotheonella” symbionts in *Theonella swinhoei:* Mixotrophy, anaerobic adaptation, resilience, and interaction. Frontiers in Microbiology 7: 1333.

Liu, L., Liu, Y., Zhang, P., Song, G., Hui, R., Wang, Z., and Wang, J. 2017. Development of bacterial communities in biological soil crusts along a revegetation chronosequence in the Tengger Desert, northwest China. Biogeosciences 14(16): 3801.

Logan, N. A. L. A., and Allan, R. N. A. N. 2008. Aerobic, endospore-forming bacteria from Antarctic geothermal soils. In Microbiology of Extreme Soils (pp. 155–175). Springer, Berlin, Heidelberg.

Männistö, M. K., Rawat, S., Starovoytov, V., and Häggblom, M. M. 2012. *Granulicella arctica* sp. nov., *Granulicella mallensis* sp. nov., Granulicella tundricola sp. nov. and Granulicella sapmiensis sp. nov., novel acidobacteria from tundra soil. International Journal of Systematic and Evolutionary Microbiology 62(9): 2097–2106.

Markowitz, V.M., Chen, I.M.A., Chu, K., Szeto, E., Palaniappan, K., Pillay, M., Ratner, A., Huang, J., Pagani, I., Tringe, S. and Huntemann, M., 2014. IMG/M 4 version of the integrated metagenome comparative analysis system. Nucleic acids research, 42(D1):D568–D573.

Medve, R. J. and Sayre, W. 1994. Heavy metals in red pines, basidiomycete sporocarps and soils on bituminous strip mine spoils. Journal of the Pennsylvania Academy of Science: 131–135.

Méheust, R., Castelle, C.J., Carnevali, P.B.M., Farag, I.F., He, C., Chen, L.X., Amano, Y., Hug, L.A. and Banfield, J.F. 2019. Aquatic Elusimicrobia are metabolically diverse compared to gut microbiome Elusimicrobia and some have novel nitrogenase-like gene clusters. bioRxiv: p.765248.

Mleczko, P. I. O. T. R. 2004. Mycorrhizal and saprobic macrofungi of two zinc wastes in southern Poland. Acta Biol. Cracov. Ser Bot 46: 25–38.

Mohagheghi, A., Grohmann, K. M. M. H., Himmel, M., Leighton, L., and Updegraff, D. M. 1986. Isolation and characterization of *Acidothermus cellulolyticus* gen. nov., sp. nov., a new genus of thermophilic, acidophilic, cellulolytic bacteria. International Journal of Systematic and Evolutionary Microbiology 36(3): 435–443.

Moyersoen, B., and Beever, R. E. 2004. Abundance and characteristics of *Pisolithus* ectomycorrhizas in New Zealand geothermal areas. Mycologia 96(6): 1225–1232.

Mrak, T., Kühdorf, K., Grebenc, T., Štraus, I., Münzenberger, B., and Kraigher, H. 2017. *Scleroderma areolatum* ectomycorrhiza on *Fagus sylvatica* L. Mycorrhiza 27(3): 283–293.

Nam, J. H., Ventura, J. S., Yeom, I. T., Lee, Y., and Jahng, D. 2016. Structural and kinetic characteristics of 1, 4-dioxane-degrading bacterial consortia containing the phylum TM7. J Microbiol Biotechnol 26: 1951-64.

Neef, A., Amann, R., Schlesner, H., and Schleifer, K. H. 1998. Monitoring a widespread bacterial group: in situ detection of planctomycetes with 16S rRNA-targeted probes. Microbiology 144(12): 3257–3266.

Ningsih, F., Yokota, A., Sakai, Y., Nanatani, K., Yabe, S., Oetari, A., and Sjamsuridzal, W. 2019. *Gandjariella thermophila* gen. nov., sp. nov., a new member of the family Pseudonocardiaceae, isolated from forest soil in a geothermal area. International Journal of Systematic and Evolutionary Microbiology 69(10): 3080–3086.

Ogo, S., Yamanaka, T., Akama, K., Ota, Y., Tahara, K., Nagakura, J., Kinoshita, A. and Yamaji, K. 2017. Growth and uptake of caesium, rubidium, and potassium by ectomycorrhizal and saprotrophic fungi grown on either ammonium or nitrate as the N source. Mycological Progress 16(8): 801–809.

Pascual, J., Wüst, P. K., Geppert, A., Foesel, B. U., Huber, K. J., and Overmann, J. 2015. Novel isolates double the number of chemotrophic species and allow the first description of higher taxa in Acidobacteria subdivision 4. Systematic and Applied Microbiology 38(8): 534–544.

Pent, M., Hiltunen, M., Põldmaa, K., Furneaux, B., Hildebrand, F., Johannesson, H., Ryberg, M. and Bahram, M. 2018. Host genetic variation strongly influences the microbiome structure and function in fungal fruiting-bodies. Environmental Microbiology 20(5):1641–1650.

Pent, M., Bahram, M. and Põldmaa, K., 2020. Fruitbody chemistry underlies the structure of endofungal bacterial communities across fungal guilds and phylogenetic groups. The ISME journal, 14(8), pp.2131–2141.

Prakash, O., Green, S.J., Jasrotia, P., Overholt, W.A., Canion, A., Watson, D.B., Brooks, S.C. and Kostka, J.E. 2012. Rhodanobacter denitrificans sp. nov., isolated from nitrate-rich zones of a contaminated aquifer. International Journal of Systematic and Evolutionary Microbiology 62(10): 2457–2462.

Quandt, C. A., Kohler, A., Hesse, C. N., Sharpton, T. J., Martin, F., and Spatafora, J. W. 2015. Metagenome sequence of *Elaphomyces granulatus* from sporocarp tissue reveals Ascomycota ectomycorrhizal fingerprints of genome expansion and a Proteobacteria-rich microbiome. Environmental Microbiology 17(8): 2952–2968. doi:10.1111/1462-2920.12840

Rinta-Kanto, J. M., Pehkonen, K., Sinkko, H., Tamminen, M. V., and Timonen, S. 2018. Archaea are prominent members of the prokaryotic communities colonizing common forest mushrooms. Canadian Journal of Microbiology 64(10): 716–726. doi:10.1139/cjm-2018-0035

Roccotiello, E., Marescotti, P., Di Piazza, S., Cecchi, G., Mariotti, M. G., and Zotti, M. 2015. Biodiversity in metal-contaminated sites–problem and perspective–a case study. In Biodiversity in Ecosystems-Linking Structure and Function. IntechOpen.

Romano, I., Poli, A., Lama, L., Gambacorta, A., and Nicolaus, B. 2005. *Geobacillus thermoleovorans* subsp. stromboliensis subsp. nov., isolated from the geothermal volcanic environment. The Journal of General and Applied Microbiology 51(3): 183–189.

Rudawska, M., Leski, T., and Stasińska, M. 2011. Species and functional diversity of ectomycorrhizal fungal communities on Scots pine (*Pinus sylvestris* L.) trees on three different sites. Annals of Forest Science 68(1): 5–15.

Schlaeppi, K. and Bulgarelli, D. 2015. The plant microbiome at work. Molecular Plant-Microbe Interactions 28(3): 212–217.

Selenska-Pobell, S., Flemming, K., Tzvetkova, T., Raff, J., Schnorpfeil, M., and Geißler, A. 2002. Bacterial communities in uranium mining waste piles and their interaction with heavy metals. In Uranium in the Aquatic Environment (pp. 455–464). Springer, Berlin, Heidelberg.

Severino, R., Froufe, H. J., Barroso, C., Albuquerque, L., Lobo-da-Cunha, A., da Costa, M. S., and Egas, C. 2019. High-quality draft genome sequence of *Gaiella occulta* isolated from a 150 meter deep mineral water borehole and comparison with the genome sequences of other deep-branching lineages of the phylum Actinobacteria. MicrobiologyOpen, 8(9), e00840.

Shi, L., Xue, J., Liu, B., Dong, P., Wen, Z., Shen, Z. and Chen, Y. 2018. Hydrogen ions and organic acids secreted by ectomycorrhizal fungi, Pisolithus sp1, are involved in the efficient removal of hexavalent chromium from waste water. Ecotoxicology and Environmental Safety 161: 430-436.

Silva, R. F., Lupatini, M., Trindade, L., Antoniolli, Z. I., Steffen, R. B., and Andreazza, R. 2013. Copper resistance of different ectomycorrhizal fungi such as *Pisolithus microcarpus*, *Pisolithus* sp., Scleroderma sp. and Suillus sp. Brazilian Journal of Microbiology 442: 613–622.

Sivakala, K. K., Jose, P. A., Thinesh, T., Anandham, R., Sivakumar, N., and Jebakumar, S. R. D. 2017. Metagenomic analysis of microbial heterogeneity and stress response Mechanisms in Desert. Canadian Journal of Biotechnology, 1(Special): 135.

Sofia, H. J., Chen, G., Hetzler, B. G., Reyes-Spindola, J. F., and Miller, N. E. 2001. Radical SAM, a novel protein superfamily linking unresolved steps in familiar biosynthetic pathways with radical mechanisms: functional characterization using new analysis and information visualization methods. Nucleic Acids Research 29(5): 1097–1106.

Soo, R.M., Wood, S.A., Grzymski, J.J., McDonald, I.R. and Cary, S.C. 2009. Microbial biodiversity of thermophilic communities in hot mineral soils of Tramway Ridge, Mount Erebus, Antarctica. Environmental Microbiology 11(3): 715–728

Szymańska, K., and Strumińska-Parulska, D. 2020. Atmospheric fallout impact on 210 Po and 210 Pb content in wild growing mushrooms. Environmental Science and Pollution Research: 1–7.

Tillakarathna, K.N., 2016. Eubacterial Endosymbionts of the Fungus Pisolithus Tinctorius (Doctoral dissertation, California State University, East Bay).

Torres-Cortes, G., Ghignone, S., Bonfante, P., and Schussler, A. 2015. Mosaic genome of endobacteria in arbuscular mycorrhizal fungi: Transkingdom gene transfer in an ancient mycoplasma-fungus association. Proceedings of the National Academy of Sciences USA 112(25): 7785–7790. doi:10.1073/pnas.1501540112

Turner, T.R., James, E.K. and Poole, P.S. 2013. The plant microbiome. Genome Biology, 14(6), p.209.

Van Den Heuvel, R. N., Van Der Biezen, E., Jetten, M. S. M., Hefting, M. M., and Kartal, B. 2010. Denitrification at pH 4 by a soil-derived *Rhodanobacte*r-dominated community. Environmental Microbiology 12(12): 3264–3271.

Wagner, I. D., and Wiegel, J. 2008. Diversity of thermophilic anaerobes. Annals of the New York Academy of Sciences 1125(1): 1–43.

Wang, H.F., Zhang, Y.G., Chen, J.Y., Guo, J.W., Li, L., Hozzein, W.N., Zhang, Y.M., Wadaan, M.A. and Li, W.J. 2015. *Frigoribacterium endophyticum* sp. nov., an endophytic actinobacterium isolated from the root of *Anabasis elatior* (CA Mey.) Schischk. International Journal of Systematic and Evolutionary Microbiology 65(4): 1207–1212.

White, T. J., Bruns, T., Lee, S. J. W. T., & Taylor, J. 1990. Amplification and direct sequencing of fungal ribosomal RNA genes for phylogenetics. PCR Protocols: A Guide to Methods and Applications 18(1): 315–322.

Wilson, A.W., Binder, M. and Hibbett, D.S., 2012. Diversity and evolution of ectomycorrhizal host associations in the Sclerodermatineae (Boletales, Basidiomycota). New Phytologist 194(4): 1079–1095.

Wilson, M.C., Mori, T., Rückert, C., Uria, A.R., Helf, M.J., Takada, K., Gernert, C., Steffens, U.A., Heycke, N., Schmitt, S. and Rinke, C. 2014. An environmental bacterial taxon with a large and distinct metabolic repertoire. Nature 506(7486): 58–62.

Wiseschart, A., Mhuantong, W., Tangphatsornruang, S., Chantasingh, D., and Pootanakit, K. 2019. Shotgun metagenomic sequencing from Manao-Pee cave, Thailand, reveals insight into the microbial community structure and its metabolic potential. BMC Microbiology 19(1): 144.

Yilmaz, P., Parfrey, L.W., Yarza, P., Gerken, J., Pruesse, E., Quast, C., Schweer, T., Peplies, J., Ludwig, W. and Glöckner, F.O. 2014. The SILVA and “all-species living tree project (LTP)” taxonomic frameworks. Nucleic Acids Research 42(D1): D643–D648.

Youssef, N. H., Farag, I. F., Rinke, C., Hallam, S. J., Woyke, T., and Elshahed, M. S. 2015. *In silico* analysis of the metabolic potential and niche specialization of candidate phylum “Latescibacteria” (WS3). PLoS One, 10(6): e0127499. doi:10.1371/journal.pone.0127499.

Yu, F., Liang, J. F., Song, J., Wang, S. K., and Lu, J. K. 2020. Bacterial Community Selection of *Russula griseocarnosa* Mycosphere Soil. Frontiers in Microbiology 11: 347.

Yun, Y., Wang, H., Man, B., Xiang, X., Zhou, J., Qiu, X., Duan, Y. and Engel, A.S. 2016. The relationship between pH and bacterial communities in a single karst ecosystem and its implication for soil acidification. Frontiers in Microbiology 7:1955.

Xia, Y., Min, H., Rao, G., Lv, Z. M., Liu, J., Ye, Y. F., and Duan, X. J. 2005. Isolation and characterization of phenanthrene-degrading *Sphingomonas paucimobilis* strain ZX4. Biodegradation 16(5): 393–402.

Zagriadskaia, Y. A., Lysak, L. V., Sidorova, II, Aleksandrova, A. V., and Voronina, E. Y. 2013. Bacterial complexes of the fruiting bodies and hyphosphere of certain basidiomycetes. Biology Bulletin 40(4): 358–364.

Zhang, H., Sekiguchi, Y., Hanada, S., Hugenholtz, P., Kim, H., Kamagata, Y., and Nakamura, K. 2003. Gemmatimonas aurantiaca gen. nov., sp. nov., a Gram-negative, aerobic, polyphosphate-accumulating micro-organism, the first cultured representative of the new bacterial phylum Gemmatimonadetes phyl. nov. International Journal of Systematic and Evolutionary Microbiology 53(4): 1155–1163.

Zhang, C.F., Ai, M.J., Wang, J.X., Liu, S.W., Zhao, L.L., Su, J., Sun, C.H., Yu, L.Y. and Zhang, Y.Q. 2016. *Herbihabitans rhizosphaerae* gen. nov., sp. nov., a member of the family Pseudonocardiaceae isolated from rhizosphere soil of the herb Limonium sinense (Girard). International Journal of Systematic and Evolutionary Microbiology 66(10): 4156–4161.

Zhang, Z., Cao, H., Song, N., Zhang, L., Cao, Y., and Tai, J. 2020. Long-term hexavalent chromium exposure facilitates colorectal cancer in mice associated with changes in gut microbiota composition. Food and Chemical Toxicology 111237.

